# Compact deep neural network models of visual cortex

**DOI:** 10.1101/2023.11.22.568315

**Authors:** Benjamin R. Cowley, Patricia L. Stan, Jonathan W. Pillow, Matthew A. Smith

## Abstract

A powerful approach to understanding the computations carried out in visual cortex is to develop models that predict neural responses to arbitrary images. Deep neural network (DNN) models have worked remarkably well at predicting neural responses [1, 2, 3], yet their underlying computations remain buried in millions of parameters. Have we simply replaced one complicated system *in vivo* with another *in silico*? Here, we train a data-driven deep ensemble model that predicts macaque V4 responses ∼50% more accurately than currently-used task-driven DNN models. We then compress this deep ensemble to identify *compact* models that have 5,000x fewer parameters yet equivalent accuracy as the deep ensemble. We verified that the stimulus preferences of the compact models matched those of the real V4 neurons by measuring V4 responses to both ‘maximizing’ and adversarial images generated using compact models. We then analyzed the inner workings of the compact models and discovered a common circuit motif: Compact models share a similar set of filters in early stages of processing but then specialize by heavily consolidating this shared representation with a precise readout. This suggests that a V4 neuron’s stimulus preference is determined entirely by its consolidation step. To demonstrate this, we investigated the compression step of a dot-detecting compact model and found a set of simple computations that may be carried out by dot-selective V4 neurons. Overall, our work demonstrates that the DNN models currently used in computational neuroscience are needlessly large; our approach provides a new way forward for obtaining explainable, high-accuracy models of visual cortical neurons.

## Main

One of the most influential computational models of sensory processing is the simple yet effective linear-nonlinear (LN) model, which uses a single spatiotemporal filter to describe a neuron’s stimulus selectivity [4, 5, 6, 7]. LN models accurately predict responses of neurons in retina and primary visual cortex (V1), and their filter parameters are easily interpretable [8, 9, 10]. However, LN models fail to predict responses from neurons in higher-order visual cortex, such as V4 and IT [11, 12], making it clear that more complicated models are needed [13, 14, 15]. Recent work has shown that after training a DNN model to perform object recognition, the DNN’s internal representations are predictive of V4 and IT responses both in human and non-human primates [1, 2, 16]. However, these task-driven DNN models have tens of millions of parameters, making it next to impossible to understand the step-by-step computations between image and response [17, 18]. Are such large DNN models necessary? In this work, we seek to determine whether models with far fewer parameters can accurately predict neural responses. If the smallest model still requires millions of parameters to be predictive, one may give up on understanding local circuit-level interactions of this “black box” and instead focus on global, layer-level operations [2, 3]. However, if the smallest model more closely resembles the LN model, we may yet have a chance to understand the step-by-step computations of visual cortical processing by analyzing a small number of simple filters [19], such as edge and curve detectors [20, 21], and basic computations, such as surround suppression [22, 23], winner-takes-all competition [24], and divisive normalization [25]. In this paper, we find the latter: We build simple yet powerful models of V4 responses to natural images and then dissect these models to understand the circuit-level computations of V4 neurons.

### Identifying compact deep neural network models of V4 neurons

We focused our efforts on predicting neural responses in macaque V4, a mid-level visual processing area whose neurons are selective for a wide range of visual features, such as edges, curves, textures, and colors [26, 27]. This diversity in stimulus preferences makes it difficult for any one model to dependably predict V4 responses across different stimulus types [28, 29]. To date, the most predictive models of V4 neurons rely on the intermediate stages of large, task-driven DNN models trained to perform object recognition [1, 30]. While probing the filters in such models has led to some success in identifying key computations [31, 32, 33], we were inspired by recent advances in model compression techniques that identified compressed DNN models with vastly fewer parameters but equivalent accuracy in the setting of object recognition [34, 35, 36]. If sufficiently small models are predictive of V4 responses, we may be able to uncover the inner computations of the compressed models. We took a two-step approach. First, because task-driven models are not directly optimized to predict V4 responses, we chose to optimize a data-driven DNN model trained on V4 responses to many images with the goal of achieving high prediction power specifically for the V4 neurons we record. Second, we designed a model compression framework to identify compressed DNN models of our large data-driven DNN model.

We built a large, data-driven DNN model that passed an input image through a task-driven DNN to obtain activity features from an intermediate layer (Fig. 1**a**, blue DNN); we then passed these features as input into an ensemble of convolutional DNNs (Fig. 1**a**, green DNNs). We used this deep ensemble to overcome overfitting: Each DNN in the ensemble overfits differently to the small amount of training data, and averaging over the ensemble averages away differences in overfitting [37]. The parameters of the deep ensemble were trained, while the task-driven model’s parameters remained fixed. Our training data comprised 44 recording sessions from 3 monkeys; each session had ∼50 neurons and∼2,000 unique images, leading to a combined total of∼78,000 unique images (see Methods). We treated each session has having a new set of neurons by assuming a new linear readout for each session (see Ext. Data Fig. 1 for detailed model diagrams); we held out 4 recording sessions (at least 1 session from each animal) for evaluation.

**Figure 1:**
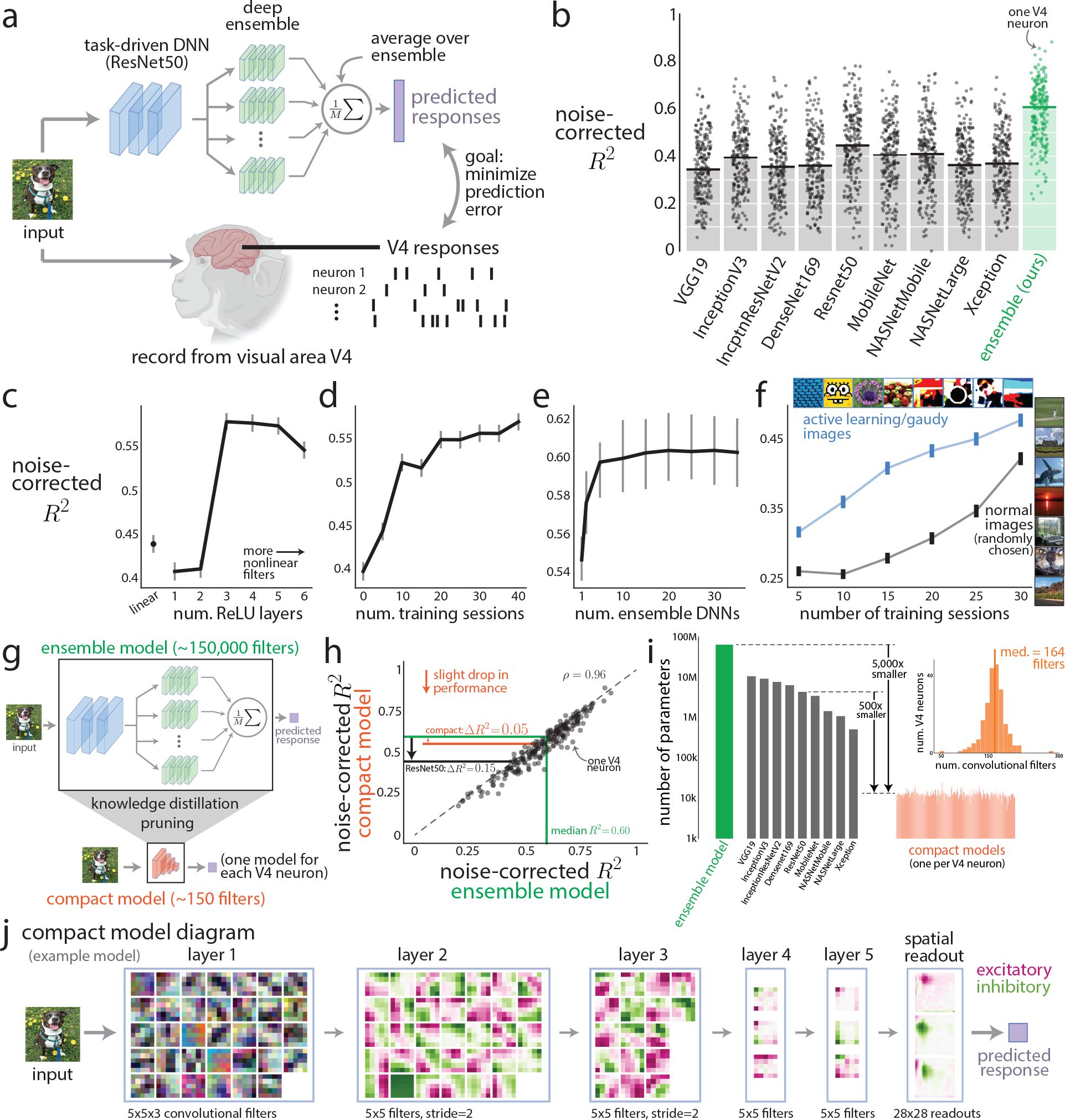
Identifying compact models of macaque V4 neurons. **a**. We presented natural images while recording from neurons in visual cortical area V4. We model the mapping between images and repeat-averaged V4 responses (spike counts taken in 100 ms bins) with a two-stage model. The first stage is to pass the image through the early and middle layers of a task-driven DNN trained to perform object recognition (blue, ResNet50). We then take the output activity of a middle layer of ResNet50 as input to the second stage—an ensemble of convolutional DNNs (green). Each ensemble DNN has the same architecture but different random initializations. The weights of the deep ensemble are shared across recording sessions, but we assume a new set of neurons each session by fitting new linear mappings between the deep ensemble’s output and V4 responses for each session. The final predicted V4 responses are the outputs of these linear mappings averaged across the ensemble (see Ext. Data Fig. 1 for detailed diagrams). **b**. Comparison of prediction performance for different DNN models on held-out images and recording sessions. DNN models included task-driven models (black) as well as our proposed data-driven deep ensemble model (green). We report noise-corrected *R*^2^, which accounts for repeat-to-repeat noise in the estimates of the repeat-averaged responses (Ext. Data Fig. 2). Each dot denotes one V4 neuron. **c**-**f**. Four modeling improvements that boosted prediction performance. These included placing a nonlinear mapping between ResNet50 features and V4 responses (**c**), training on a large number of recording sessions (**d**), using a deep ensemble with many small DNNs (**e**), and training on images chosen adaptively by active learning and gaudy images versus randomly-chosen normal images (**f**). See Methods for details on each analysis. Lines denote medians, error bars denote 90% bootstrapped confidence intervals. **g**. Framework to identify compact models. We take our large deep ensemble model (top) and use the model compression techniques (knowledge distillation and pruning) to identify a compact model, one for each V4 neuron (bottom). **h**. Prediction performance on held-out V4 responses for the deep ensemble model versus that of the compact models. Each dot denotes one V4 neuron; the dashed line denotes the same level of prediction. On average, the compact models predicted V4 responses only slightly worse than the deep ensemble model with a decrease in median noise-corrected ∆*R*^2^ = 0.05 (orange line), much smaller than that of task-driven ResNet50 features (∆*R*^2^ = 0.15, black line). The *R*^2^s between compact and ensemble model predictions across neurons were highly correlated (*ρ* = 0.96), suggesting that no group of V4 neurons (e.g., “dot detectors”) was more poorly explained than another group by the compact models. **i**. Number of parameters (including linear mappings) for our deep ensemble model (green), task-driven models (black), and compact models (orange). The *y*-axis is in log-scale. Inset: Total number of convolutional filters for each compact model across V4 neurons; the median was 164 filters, whereas our deep ensemble model had∼150,000 filters. **j**. Diagram for an example compact model showing all weight parameters for its convolutional filters. Pink denotes positive (or excitatory) weights, and green denotes negative (or inhibitory) weights. Because layer 1 filters directly take the RGB image as input, the weights are colored based on RGB channels. For clarity, the mixing weights and batchnorm weights are not shown (see Ext. Data Fig. 1).

We found that prediction performances of task-driven DNN models (Fig. 1**b**, black,∼40% explained variance, noise corrected) were consistent with those reported in previous studies [12, 38, 30, 39] (but see [40] and Ext. Data Fig. 2). The prediction performance of our ensemble model was 50% better than that of task-driven models (Fig. 1**b**, green, ∼ 60% explained variance, noise corrected). Despite being trained only on our recorded V4 neurons, our deep ensemble model was predictive of V4 responses from a separate study (Ext. Data Fig. 2), suggesting our model generalizes well to held out V4 neurons. The substantial boost in performance was primarily the result of four modeling improvements. First, allowing for a nonlinear mapping (i.e., the deep ensemble) between task-driven DNN features (e.g., ResNet50) and V4 responses better predicted responses (Fig. 1**c**) than using a linear mapping typical of most studies [12, 30, 39, 41]. Second, training on a large number of recording sessions led to better prediction, as expected (Fig. 1**d**). Third, using a DNN ensemble versus a single DNN increased prediction performance (Fig. 1**e**). Fourth, a portion of our training images were chosen adaptively in closed-loop experiments via active learning [42, 43, 44] and synthetically-generated “gaudy images”, previously shown to improve training of DNN models [45]. Training on responses to these images led to better prediction than randomly-chosen natural images (Fig. 1**f**, blue line above black line). Together, these improvements helped to substantially boost prediction performance of the deep ensemble model by 50% over current task-driven models—this alone was a significant advance in state-of-the-art prediction of V4 responses. However, the deep ensemble model’s 60 million parameters made interpreting any internal computations next to impossible, a key challenge in making DNN models usable for understanding vision.

To overcome this challenge, we sought ways to compress the deep ensemble model as much as possible while maintaining high prediction power—how small can the model be without giving up predictive power? To address this question, we used in tandem a pair of model compression techniques known as knowledge distillation [35] and pruning [36, 46, 47, 48, 49] (Fig. 1**g**). We first performed distillation by collecting the deep ensemble model’s predicted responses to 12 million natural images; we then used these image-response pairs to train a much smaller DNN with 5 layers (100 convolutional filters per layer). Training this small DNN model directly on our real data failed due to overfitting (median noise-corrected *R*^2^ = 0.11), hence the need for distillation. We then pruned this distilled DNN by ablating convolutional filters that contributed little to the model’s output and then re-trained; we stopped pruning when prediction performance decreased by 5% relative to that of the deep ensemble model. We call the resulting pruned model a *compact* model; for each V4 neuron, we fit one compact model. The prediction performance of the compact models (Fig. 1**h**, median noise-corrected *R*^2^ = 0.55) was only slightly less than that of the ensemble model (median noise-corrected *R*^2^ = 0.60) but much greater than that of the top performing task-driven DNN model, ResNet50 (median noise-corrected *R*^2^ = 0.45). Importantly, the compact models had ∼5,000 times fewer parameters than our deep ensemble model (Fig. 1**i**, ‘ensemble’) and ∼ 500 times fewer parameters than the task-driven ResNet50 (Fig. 1**i**, ‘ResNet50’). Indeed, a compact model had few enough convolutional filters (Fig. 1**i**, inset) for all of them to be viewed in one diagram (Fig. 1**j**), similar to how the filter weights of LN models are inspected. This result indicates that the current task-driven DNN models used to predict V4 responses are substantially larger—by orders of magnitude—than needed. Moreover, our compact models are small enough ( ∼ 150 filters in total per model) for us to systematically interrogate their inner workings, which we pursue in later sections.

### Causally testing the compact models

A natural first step in understanding the computations of the compact models is to identify their preferred stimuli [40, 43, 50, 51, 52, 53, 54, 55, 56]. For example, a retinal ON cell strongly prefers a white spot surrounded by black [9, 57], and a V1 neuron strongly prefers a sinusoidal grating with a specific orientation, phase, and spatial frequency [58]. To find the preferred stimuli of each compact model, we synthesized an image that maximized that model’s output using gradient techniques (see Methods). Examples of these maximizing synthesized images revealed edges, curves, dots, and textures (Fig. 2**a**), as expected from previous studies [20, 21, 59, 60, 56]. One concern about drawing conclusions from these preferred stimuli—which resemble but are not natural images— was that we had not tested whether they actually more strongly drive the responses of real V4 neurons relative to natural images. To address this concern, we designed a set of causal experiments to test whether the preferred stimuli of our compact models matched those of real V4 neurons. Each causal experiment followed the same procedure: We first trained compact models on previous recording sessions; we then probed each model for its preferred stimulus; finally, we presented these probed images in a future session along with randomly-chosen normal images for reference. In this way, we *causally* tested the predictions of the compact models outside the distribution of randomly-selected natural images.

**Figure 2:**
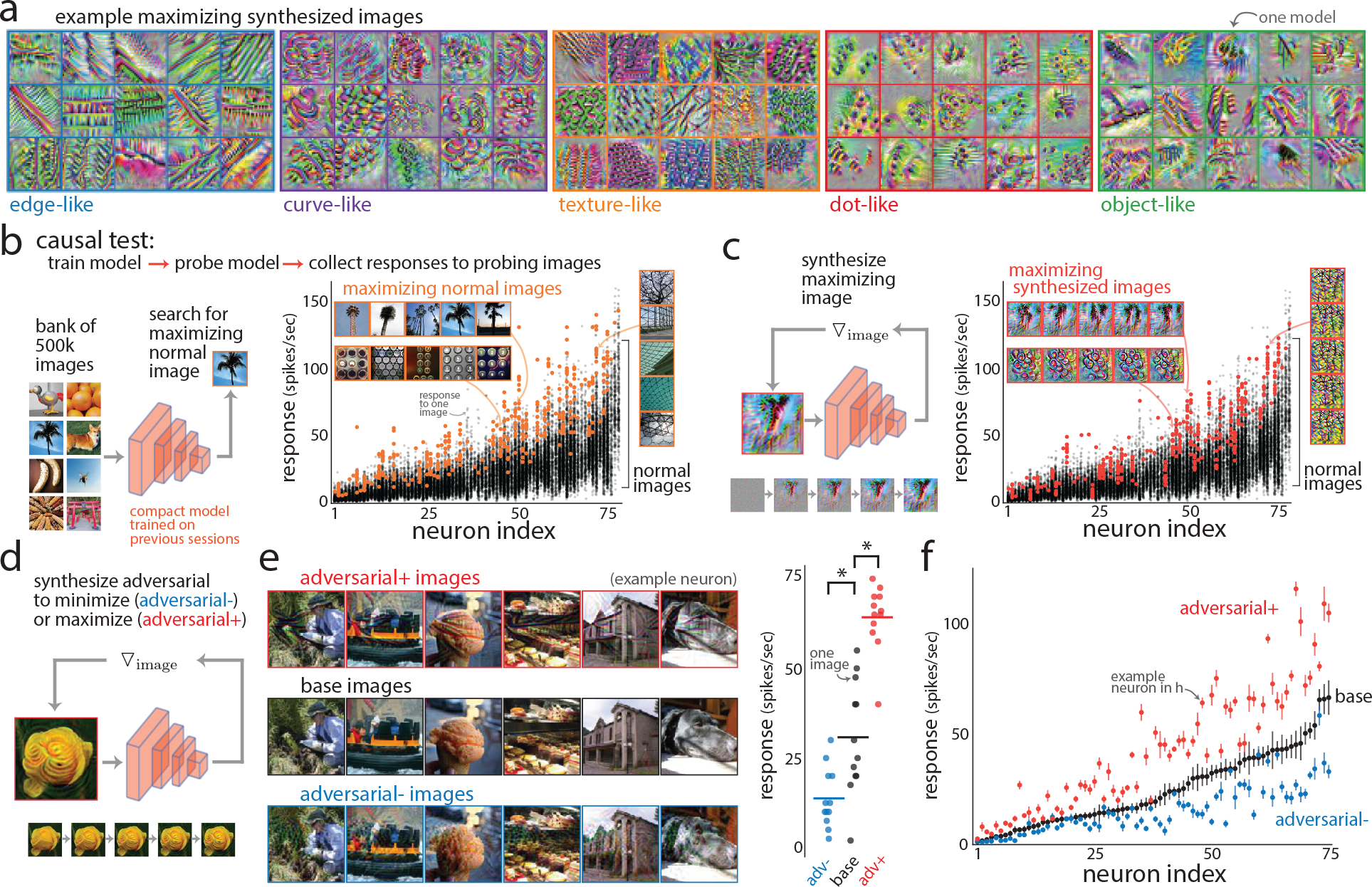
Causally testing the predictions of compact models. **a**. Preferred synthetic stimuli of selected compact models (one stimulus per model; see Ext. Data Fig. 3 for all compact models). Each image is optimized via gradient ascent to maximize a model’s output response (see Methods). For illustrative purposes, preferred stimuli were loosely placed by eye into categories (e.g., “edge-like” detectors, “curve-like” detectors, etc.). **b**. Causal test in which we trained a compact model on previous recording sessions, probed the model to identify images from a bank of 500,000 candidate images that maximize the model’s output (left), and then recorded V4 responses to these maximizing normal images in future sessions (right). The maximizing normal images drove larger V4 responses (orange dots, normalized percent change computed as 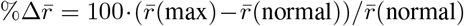: mean 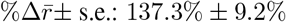, where 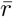 is the mean response over images) than responses to randomly-chosen normal images (black dots). Each dot denotes the repeat-averaged response to one image. Each neuron index (*x*-axis) refers to one of the 78 neurons recorded from two monkeys, ordered by average response to the normal images. Insets show maximizing normal images for three example neurons. **c**. Causal test for maximizing synthesized images of the compact models. Each synthesized image started as a white noise image and iteratively changed via gradient ascent by propagating the gradient with respect to the model’s output back through the compact model (example iterations shown in bottom left inset). In future recording sessions, the maximizing synthesized images elicited larger V4 responses (red dots, 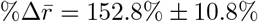) than responses to randomly-chosen normal images (black dots). Insets show the maximizing synthesized images for three example neurons (same as in **b**). **d**. Causal test for adversarial images of the compact models. We define an adversarial image as a slight perturbation to some base image that yields a large change in the model’s output. Adversarial images were synthesized via gradient descent to minimize the model’s output (‘adversarial-’) or gradient ascent to maximize the model’s output (‘adversarial+’). Synthesis stopped when differences in pixel intensities between the base image and the adversarial image passed a threshold. **e**. Example adversarial+, base, and adversarial-images for an example compact model (left). As predicted by the compact model, a corresponding real V4 neuron responded less to the adversarialim-ages (blue dots) and more to the adversarial+ images (red dots) compared to responses to the base images (black dots). Each dot denotes the repeat-averaged response to one image; lines denote means, and asterisks denote *p <* 0.05 (permutation test). **f**. In future recording sessions, adversarial+ images drove larger responses (red dots, 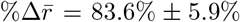) and adversarial-images drove smaller responses (blue dots, 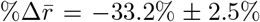) compared to responses to base images (black dots). Dots denote means, and error bars denote 1 s.e.m. across 10 or more base images per neuron.

We first probed each compact model using “preferred stimuli” selected from our large image database (Fig. 2**b**, left panel). These maximizing natural images drove V4 responses ∼ 2.4 times larger than the average V4 response to randomly-chosen images (Fig. 2**b**, orange dots above black dots). Next, we probed the compact models by synthesizing images to maximize the response of a given model neuron (Fig. 2**c**, left panel). We found that the synthesized maximizing images elicited V4 responses ∼ 2.5 times larger than the average response to a randomly-chosen image (Fig. 2**c**, red dots above black dots). Interestingly, the maximizing natural images tended to drive V4 responses as strongly as the maximizing synthesized images, even though the latter were predicted by the compact models to elicit the largest responses (Ext. Data Fig. 4). This suggests that the compact models were overly confident in predicting images outside the distribution of natural images—which may be remedied by training on out-of-distribution images [61, 62]—and that one should not rule out searching over a large number of natural images (in this case, 500,000 images) when identifying a neuron’s preferred stimulus.

We viewed the causal tests of maximizing natural and synthesized images as coarse tests: We strongly drove neurons with unnaturalistic images outside the range of their usual working regime. We wondered whether predictions of the compact models also held while not deviating far from the distribution of natural images. To this end, we slightly perturbed a natural image to yield large changes in the model’s output; in other words, we identified *adversarial* images of the compact models [63, 64]. To identify an adversarial image (Fig. 2**d**), we started with a base image and used a gradient method to either maximize (‘adversarial+’) or minimize (‘adversarial-’) the model’s output (similar to identifying the maximizing synthesized image). Once the adversarial image deviated too much from the base image (an average change of 0.04 in pixel intensity, where pixel intensity could range from 0 to 1), synthesis stopped. The resulting adversarial images were perceptually similar to the base image (Fig. 2**e**, left, ‘adversarial+’ and ‘adversarial-’ versus ‘base’) but yielded large response changes for an example V4 neuron (Fig. 2**e**, right), consistent with the compact model’s predictions. We found this to be a strong effect across V4 neurons (Fig. 2**f**). Thus, despite a practically infinite number of ways to slightly perturb an input image, the compact models correctly predicted two perturbations (‘adversarial+’ and ‘adversarial-’) that strongly drove V4 responses in opposite ways. Taking the results of these causal tests together (we also performed more causal tests, see Ext. Data Fig. 4), we conclude that the compact models accurately capture the stimulus preferences of their V4 neurons, even for stimuli that lay well outside the distribution they were trained on.

### Compact models specialize via a consolidation step

Given the compact models’ high level of accuracy in predicting held-out V4 responses and preferred stimuli, we were motivated to investigate the inner workings of the compact models to gain more insight into the computations carried out by real V4 neurons. We started by comparing the number of filters for each layer, which could vary as determined by our pruning algorithm. One possibility was that the number of filters would gradually increase from the earliest layer to the deepest layer—similar to the architectures of the most successful task-driven DNNs [65, 66, 63]—as a higher dimensionality (i.e., more filters) tends to make representations more linearly-separable [29]. An additional possibility was that the number of filters may vary greatly between compact models (where each model corresponds to one neuron), reflecting the varying complexity of models needed to explain the heterogeneity of stimulus preferences across different neurons (Fig. 2**a**). However, we found no evidence to support either scenario. Instead, we found that the early layers had large numbers of convolutional filters (Fig. 3**a**, layers 1-3) while the later layers had substantially fewer filters (Fig. 3**a**, layers 4-5 and spatial readout). This trend could not be explained by our method of pruning (Ext. Data Fig. 2). Because this steep decrease in the number of filters between layers 3 and 4 indicates the model consolidates information across many filters into only a few, we refer to this step as the “consolidation step”. That this abrupt consolidation occurred at the same layer for almost all models suggests that the consolidation step is a hallmark of V4 processing.

**Figure 3:**
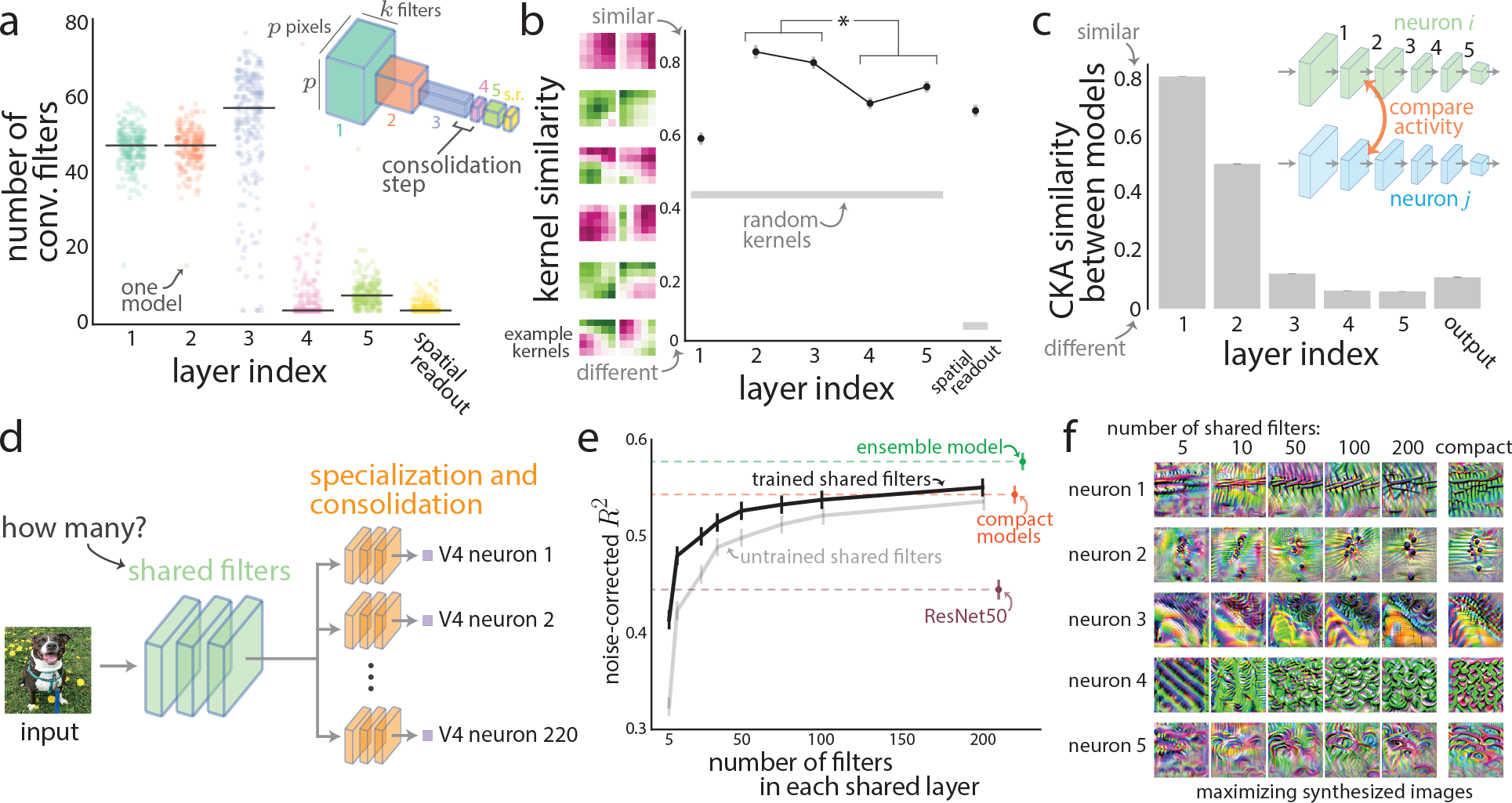
Compact models share similar filters in early layers but then heavily specialize via a consolidation step. **a**. Number of convolutional filters per layer as determined by our model compression framework; each layer potentially has up to 100 filters. Each dot denotes one compact model; lines denote medians. Layers 1 and 2 have the same number of convolutional filters as defined by the compact model’s architecture. Inset: diagram depicting activity map shapes (*p × p × k* for *p* pixels and *k* filters) for each layer. **b**. Kernel similarity between kernel weights of convolutional filters from the same layer across compact models. A kernel similarity close to 1 indicates that any convolutional filter from a given layer will have a closely matching filter (i.e., matching in its kernel weight pattern) from the same layer. For reference, we assessed the kernel similarity for filters whose weights are drawn randomly from a standard normal distribution (gray). Because layer 1 weights were not smoothed during training and the spatial readout filters were much larger (28 *×* 28 pixels vs. 5 *×* 5 pixels), it was not fair to compare the kernel similarities of these layers to the other remaining layers (layers 2-4). Layers 2 and 3 had a significantly higher mean kernel similarity than that of layers 4 and 5 (*p <* 0.002, permutation test). Dots denote mean, and error bars denote 1 s.d. across 500 runs of sampling pairs of models (see Methods). **c**. Centered kernel alignment (CKA) similarity between the activity of the same layer for two compact models. The activity is the output of each layer’s filters for 10,000 normal images. A CKA similarity close to 1 indicates that the two layers have near identicial representations up to a rotation. Layers 1 and 2 had significantly higher CKA similarity than that of layers 3 and 4 (*p <* 0.002, permutation test). The error bars denoting 1 s.e.m. are small due to the large number of pairs of models ( ∼ 23,000 pairs). **d**. Diagram of a ‘shared’ compact model to predict all V4 neurons together. The model constrains each V4 neuron to use the same shared filters in the first 3 layers (i.e., early layers), while the remaining 3 layers allow for consolidation and specialization (see Methods). It is unknown how many shared filters in the early layers are needed to explain V4 responses. **e**. Prediction performance of V4 responses while varying the number of shared filters in the early layers. For each number, a new compact model was trained via distillation by using the responses of the ensemble model (‘trained shared filters’); pruning was not performed. Fixing the kernel weights of the early layers to their initial random values (but allowing all other parameters in the early layers to be trained) led to worse prediction performance (gray line below black line). For reference, we re-computed performance for the ResNet50 features, the compact models, and the deep ensemble model (bottom, middle, and top dashed lines, respectively). Dots denote means; error bars denote 1 s.e.m. **f**. Maximizing synthesized images of the shared compact model with different numbers of shared filters (columns) as well as individual compact models (rightmost column) for five example V4 neurons.

Why would such a consolidtion step be needed? We reasoned that the early layers may represent a basis set of filters that provide a rich set of features useful for downstream neurons to specialize, much like how a bank of Gabor filters captures edge information useful for a variety of vision tasks [67, 68, 69, 70]. If true, we would expect to see that the filters and representations of the early layers are similar. We performed a coarse test of filter similarity by comparing the kernel weights of convolutional filters within the same layer across models and found that the filter patterns of early layers were more similar to one another than those of later layers (Fig. 3**b**). We then performed a more fine-grained test for similarity by comparing the internal representations of each layer between models. We chose the Centered Kernel Alignment (CKA) as our similarity metric, commonly used to compare the inner representations of two deep neural networks [71]. CKA similarity assesses to what extent two representations—in our case, the activity maps of 10,000 images from the same layer for two compact models—match in correlation between (rotated) linear dimensions and in eigenspectra (see Methods). We found that the representations of the early layers were substantially more similar than those of later layers (Fig. 3**c**). Indeed, the CKA similarity of outputs between any two models was low (Fig. 3**c**, ‘output’; the average signal correlation squared between models was *ρ*^2^ = 0.11); when controlling for overlap in spatial receptive fields, the CKA similarity was even lower (Fig. 3**c**, layer ‘5’ vs. ‘output’). This suggests that each V4 neuron specializes in a unique way and explains the diversity in stimulus preferences.

From these results arises the following picture: Each compact model reads from a shared basis set of filters extracting low-level features and then specializes by heavily consolidating these features. Based on this picture, only a small number of filters (e.g., less than 100) in the early layers would be needed to explain the activity of all recorded V4 neurons. This is surprising, as presumably there are hundreds to thousands of V1 and V2 neurons with overlapping spatial receptive fields but different Gabor-like filter patterns [68, 72]. To test this, we built a “shared” compact model in which the filters of first 3 layers were shared to predict all of our held out V4 neurons (Fig. 3**d**, green) versus one compact model per neuron as before. We trained the model on the deep ensemble responses via distillation (see Methods). We found that as few as 10 filters per shared layer (i.e., 30 filters total in layers 1-3) were needed to surpass the prediction performance of linearly mapping ResNet50 features (Fig. 3**e**, black line above the bottom dashed line). Prediction performance began to plateau at around 50 shared filters (Fig. 3**e**, black line); we further confirmed that the maximizing synthesized stimuli for 50 shared filters were similar to those for 100 or 200 shared filters (Fig. 3**f**). Fixing the weights of the shared convolutional filters to their initial random values (but allowing all other parameters, including mixing weights, to be trained, see Methods) yielded worse prediction performance than training these weights (Fig. 3**e**, line for ‘untrained shared filters’ below line for ‘trained shared filters’), ruling out the possiblity that any set of basis filters may achieve high prediction. We conclude that the diversity in stimulus preferences observed across V4 neurons occurs despite sharing a small number of filter types in the early stages. In other words, diverse specialization in V4 tuning can be achieved by reading out from a small bank of Gabor-like filters (e.g., small populations of V1 and V2 neurons)—each V4 neuron (or class of V4 neurons) does not need its own dedicated V1 and V2 neural circuit. The diverse specialization of V4 neurons likely arises during the consolidation step, whose precise readouts vastly differ between V4 neurons. Thus, understanding the computations occuring in the consolidation step is key to understanding the response profile of a V4 neuron.

### The computations of a V4 dot detector

To demonstrate that we can understand a V4 neuron’s computations by focusing on its consolidation step, we chose to investigate an exemplar compact model with a salient specialization: dot detection. This compact model’s preferred stimuli greatly emphasized dots (Fig. 4**a**). We further probed this model by presenting artifical dot stimuli that varied in location, size, and number; the model was highly selective to 3-4 small dots to the right of the image (Fig. 4**b** and Ext. Data Fig. 5) and tended to be invariant to dot positions within its receptive field (Fig. 4**a**). This preference was not an artifact of the model: We verified the existence of real V4 neurons that resembled dot detectors via causal tests (Ext. Data Fig. 6). Although the presence of dot detectors in V4 is interesting in its own right (e.g., representing a bias in the visual system toward detecting facial features such as eyes or arrays of faraway objects such as a flock of geese [56]), we were more interested in the computations that occur between the retinal inputs and the responses of a V4 dot detector. Thus, we investigated the inner computations of our chosen compact model—which maps images to V4 responses—to understand how a neural circuit may implement a dot detector.

**Figure 4:**
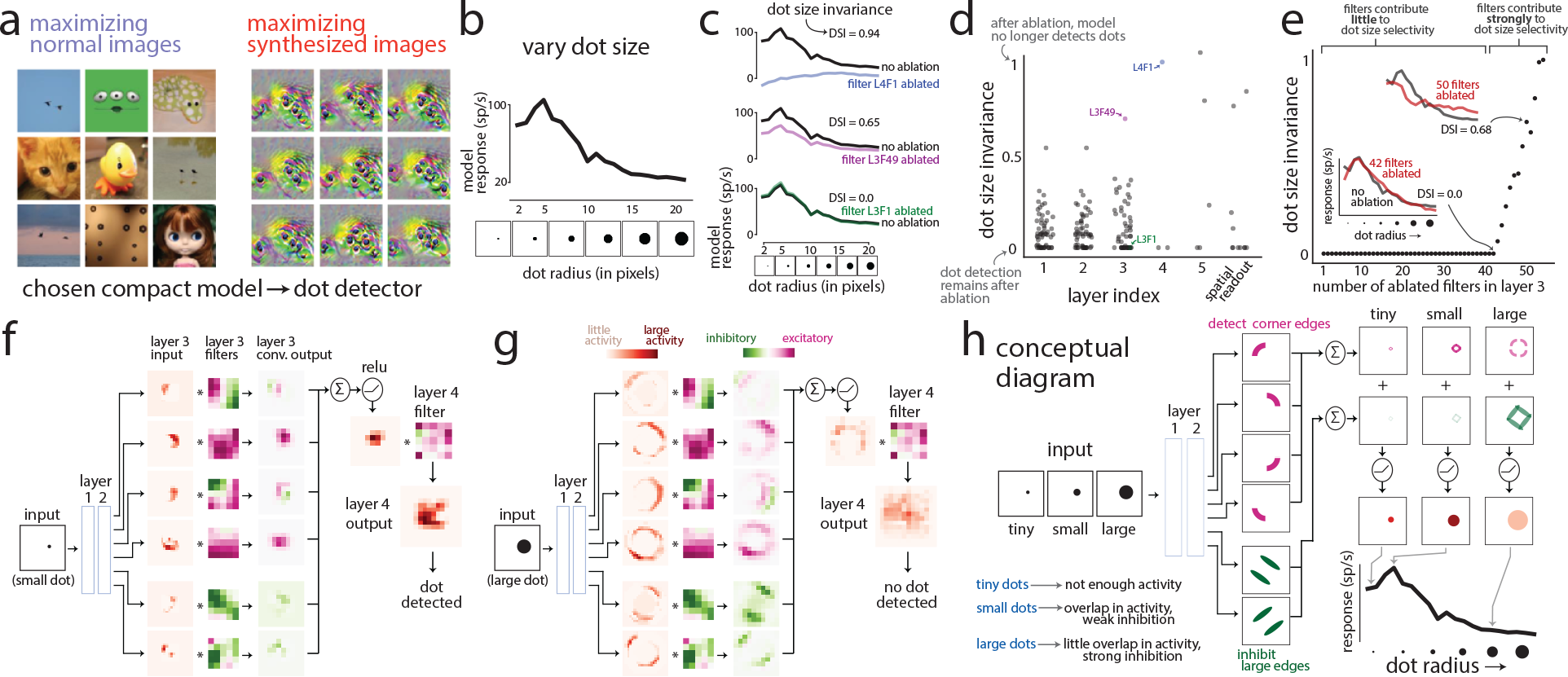
Uncovering the internal computations of a dot-detecting compact model. **a**. Preferred stimuli (maximizing normal and synthesized images) for a compact model chosen for its resemblance to a dot detector. These preferred stimuli drove large model responses well beyond the response range for normal images. **b**. For the chosen compact model in **a**, model responses to artificial dot stimuli in which we varied the size of a dot centered in the preferred location of the model. The compact model was also selective to dot location and dot number (Ext. Data Fig. 5). **c**. Responses from the model for either no ablation (black lines) or ablated filters (color lines) to artificial stimuli varying in dot size. An ablated filter (‘L4F1’ stands for Layer 4, Filter 1) has its kernel weights set to 0. We measured each filter’s dot size invariance (DSI) as the extent to which responses change due to ablation (i.e., the change between black and color lines, see Methods). A DSI = 1 indicates that, after filter ablation, the model is invariant to dot size (top panel, blue line is flat)—in other words, that filter is necessary for the model’s dot selectivity. **d**. Dot size invariance for each individual convolutional filter in the chosen compact model. Color dots correspond to example filters in **c. e**. Cumulatively ablating filters in layer 3 to identify the subset of filters that strongly contribute to dot selectivity. To do this, we keep ablating filters until DSI = 1 (i.e., until we observe a substantial change in the model’s responses). For each iteration, we choose a filter from the remaining filters that, once ablated, increases DSI the *least*. Little to no increase in DSI indicates that the ablated filter weakly contributes to dot selectivity (leftmost dots); a large increase indicates a strong contribution to dot selectivity (right-most dots). **f**. Investigating the layer 3 filters that strongly contribute to dot selectivity (i.e., 6 of the 12 rightmost dots in **e**). We pass as input an image with a small dot at the compact model’s preferred location (‘small dot’). The input activity map for each layer 3 filter (‘layer 3 input’) represents the processed image after the first two layers; each activity map is then convolved by its corresponding layer 3 filter (‘layer 3 filters’) to produce the convolved output (‘layer 3 conv. output’). This convolved activity is then summed element-wise across filters, passed through a ReLU, and then convolved again by a layer 4 filter to produce the processed output activity (‘layer 4 output’). In this case of a small dot, the output activity is large (dark red), indicating the presence of a dot (‘dot detected’). For clarity, the activity maps were cropped around the dot’s location; activity outside this cropped region was a constant value. **g**. Analyzing the same filters as in **f** except for an input image with a large dot (‘large dot’). In this case, the output activity (‘layer 4 output’) is small (light red), indicating that no dot is present (‘no dot detected’). **h**. Illustrative diagram describing the computations in the dot detector compact model. The 4 excitatory filters (pink) detect the 4 corner edges of the dot while the 2 inhibitory neurons (green) detect large edges. For a tiny dot (‘tiny’), the excitatory activity is weak, leading to a weak response. For a small dot (‘small’), the strong excitatory activity overlaps when summed (producing an even larger response) while inhibition is weak, leading to an overall large response. For a large dot (‘large’), the excitatory activity is strong but does not overlap; in addition, inhibition is strong. This leads to an overall weak response.

One might expect that a dot detector employs filters that directly search for dots of various sizes. We would thus expect to find convolutional kernels with center-surround receptive fields, but this was not the case for our example dot detector (Ext. Data Fig. 5). Where, then, does dot size selectivity arise? To answer this, we devised a metric called the dot size invariance (DSI) index that measures the extent to which each filter contributes to dot size selectivity. A DSI close to 1 indicates that when a selected filter is ablated (i.e., its weights are set to 0), the compact model becomes invariant to dot size—in other words, the intact filter strongly contributes to dot size selectivity (Fig. 4**c**, ‘filter L4F1 ablated’). On the other hand, a DSI close to 0 indicates that after ablating a selected filter, the model remains selective for dot size with little change to its output (Fig. 4**c**, ‘filter L3F1 ablated’). We found that almost all filters in the early layers weakly contributed to dot size selectivity (Fig. 4**d**, layers 1-3, DSIs *<* 0.5). Interestingly, a filter in layer 4 strongly contributed to dot selectivity (Fig. 4**d**, ‘L4F1’, DSI *≈* 1), even though no single layer 3 filter was a strong contributor to dot selectivity (Fig. 4**d**, layer 3, low DSI values). This suggests that dot detection emerges after the consolidation step by integrating information from multiple layer 3 filters.

To isolate which layer 3 filters most contributed to dot size selectivity, we performed a cumulative ablation procedure in which we chose one of the remaining filters that, once ablated, led to the smallest increase in DSI (i.e., the filter that contributed *least* to dot size selectivity). This chosen filter stayed ablated for the rest of the procedure, and in a greedy manner we proceeded to choose the next filter to ablate from the remaining unablated filters. We found that ablating as many as 40 filters together out of the possible 54 filters in layer 3 still retained a low DSI (Fig. 4**e**, bottom left inset); only a group of ∼ 10 filters contributed to dot selectivity (Fig. 4**e**, rightmost dots). We note that the weak contributing filters (Fig. 4**e**, leftmost dots) likely serve other computational purposes, such as filtering background textures, inhibiting visual features other than dots, or contributing to selectivity of other features such as the number of dots (Ext. Data Fig. 5).

We further investigated the 6 most salient of the contributing filters in layer 3 (Fig. 4**e**, rightmost dots) by observing how these filters responded to dot images. For an input image of a small dot, 4 of these filters received input activity that tiled the four corner curves of the dot (Fig. 4**f**, ‘layer 3 input’ column, top 4 filters); their excitatory kernel weights led to output activity with large positive values (Fig. 4**f**, ‘layer 3 conv. output’ column, top 4 filters). The remaining 2 filters were inhibitory for large, oriented edges in their input (Fig. 4**f**, ‘layer 3 filters’ column, bottom 2 filters). In this case, the small dot did not have large edges, and the output activity of these 2 inhibitory filters was small but negative (Fig. 4**f**, ‘layer 3 conv. output’ column, bottom 2 filters). The output activity for all filters was then elementwise summed and passed through a ReLU activation function. The strong overlap in positive activity across the top 4 filters and the weak inhibition by the bottom 2 filters yielded large positive activity (Fig. 4**f**, ‘layer 4 output’) which, after summing across spatial locations, led to the result of a dot being detected.

For an input image of a large dot, the story changed. Although the top 4 filters still responded to the four corner curves of the dot, their output activity no longer spatially overlapped (Fig. 4**g**, top 4 rows), leading to weak excitation. In addition, because the large dot contains large, prominent edges, the bottom 2 filters provided strong inhibition (Fig. 4**g**, bottom 2 rows), cancelling some of the excitation from the 4 excitatory filters. The end result was a small level of output activity (compare ‘layer 4 output’ in Fig. 4**f** and **g**), leading to the result of no dot being detected. We condensed these results into a diagram of how this compact model operates as a dot detector (Fig. 4**h**); the important computations occurred in layer 3, consistent with our hypothesis that a V4 neuron specializes via its consolidation step. Although here we focused on the model’s selectivity to dot size, the model also has a preference for multiple dots (Fig. 4**a**, right panel). This preference largely stems from the spatial structure of the layer 4 filter’s kernel weights (Fig. 4**g**, ‘layer 4 filter’), which has sparse, large magnitude weights along its perimeter to promote distance between dots (see Ext. Data Fig. 5 for a thorough analysis on dot number selectivity).

## Discussion

To understand the responses of visual cortical neurons, we seek computational models that are both highly predictive *and* explainable. To this end, we built a data-driven deep ensemble model, trained on V4 responses to tens of thousands of images, that improved prediction by 50% over current state-of-the-art task-driven DNN models. We then compressed this large DNN model to obtain *compact models* that are∼ 500 times smaller than leading task-driven DNN models, yet predict V4 responses as well as our highly-predictive data-driven DNN model. The small size of each compact model ( ∼150 convolutional filters in total) allowed us to systematically interrogate the model’s inner workings. Analyzing these compact models revealed that, consistent with previous studies [28, 40, 56, 73, 74, 75], V4 neurons form a diverse set of highly-specialized feature detectors, which are seemingly needed to form a rich basis for goal-oriented object recognition [27, 29, 31, 32, 33]. We found that such diverse specialization likely arises by reading out from a surprisingly small number of early filters resembling Gabor-like V1 neurons [67, 68] and texture-preferring V2 neurons [76, 77]; dedicated, separate circuits in early visual processing are not needed for seemingly disparate types of V4 feature detectors (e.g., edge, curve, texture, and dot detectors). A key prediction from our results is that to achieve their specializations, V4 neurons form precise synaptic connections in order to properly balance the relevant input features (e.g., reading out from excitatory corner-detectors and inhibitory large edge detectors to form a dot detector); in addition, the same V1 or V2 neuron likely synaptically projects (directly or indirectly) to multiple V4 neurons to be “re-used” for different specializations. These predictions can be tested with anatomical tracing [78, 79, 80] and circuit perturbation [81, 82, 83] experiments. We only considered simple convolutional layers in our compact models, but our model compression framework is general: Adding more biologically-plausible mechanisms, such as surround suppression [22, 23], divisive normalization [25, 84], and recurrence [85], is straightforward (i.e., estimating their parameters is tractable because of the near infinite training data via distillation training) and may lead to even smaller models. Our work focuses on one aspect of vision—the encoding of visual features; an exciting future direction is to combine encoding models with models of trial-to-trial variability [86, 87, 88, 89, 90], adaptation [91], spatial attention [92, 93, 94, 95], arousal [96, 97], visual crowding [98, 99], saccade planning [100, 101, 102], perceptual learning [103], among other functions of V4 [27]. Overall, our compact models demonstrate that the task-driven computational models of the visual cortex are needlessly large. We provide a path forward to obtain highly-predictive *and* explainable models that will help guide new experiments to record, perturb, and anatomically trace visual cortical neurons, uncovering the underlying computations of higher-order visual cortical neurons.

## Acknowledgments

We thank N. Rafidi for providing comments on the manuscript. This work was supported by a C.V. Starr Fellowship to B.R.C.; an NIH grant (F31EY031975) to P.L.S.; a Simons Collaboration on the Global Brain Investigator Award (SCGB AWD543027), NIH BRAIN Initiative grants (NS104899 and R01EB026946), and a U19 NIH-NINDS BRAIN Initiative Award (5U19NS104648) to J.W.P.; and NIH grants (R01MH118929, R01EB026953, and R01EY029250) and an NSF NCS grant (1734916/1954107) to M.A.S.

## Data availability

Data including responses and stimuli from all training and test sessions will be publicly available by publication.

## Code availability

Model weights and code will be publicly available by publication.

## Competing interests

The authors declare no competing interests.

## Author contributions

B.R.C., P.L.S., J.W.P., and M.A.S. conceived of and designed the study. B.R.C. and P.L.S. designed and performed the closed-loop experiments; B.R.C. provided image stimuli, and P.L.S. recorded the electrophysiological data. B.R.C. designed, trained, and analyzed the models. B.R.C wrote the manuscript with input from P.L.S., J.W.P., and M.A.S.

## Methods

### Experimental animals

Three adult male animals (Macaca mulatta) WE, PE, and RA were used for this study. Experimental procedures were approved by the Institutional Animal Care and Use Committee of Carnegie Mellon University and were performed in accordance with the United States National Research Council’s Guide for the Care and Use of Laboratory Animals.

### Neurophysiological experiments

#### Electrophysiology

We recorded extracellular activity from populations of V4 neurons in three awake, head-fixed monkeys; details about surgical procedures and electrophysioligcal recordings have been reported in a previous study that shared two monkeys [104]. Briefly, animals were surgically implanted with a titanium headpost to allow for head fixation during experiments. We chronically implanted a 96-electrode array (Blackrock Microsystems; 1 mm in electrode length, 400 *µ*m spacing in a 10 *×* 10 grid) in the left hemisphere V4 of each animal. V4 arrays were implanted on the prelunate gyrus medial to the inferior occipital sulcus. Electrodes of 1 mm in length would likely reach layers 4 and 5 of V4, where the cortical thickness is close to 2 mm; the precise layer of our recordings was not verified because it would have required sacrificing the animals for histology.

#### Processing of V4 responses

To process the recorded spike signals, we used an automated deep learning pipeline [105] built with custom MATLAB software. This pipeline separate spike waveforms from noise on each channel. We disregarded units with firing rates less than 1 spike/sec. For each unit, we computed an unbiased estimator of the signal-to-noise ratio (SNR) across images and repeats [106]; we disregarded units with the recommended SNR less than 0.15. Our data likely consisted of both single and multi-units; we refer to each unit as a “neuron” for simplicity. Although a subset of neurons were likely the same across sessions for the same implanted array, similar to previous studies [40], we observed that after spike processing the number of neurons differed across sessions. This was expected due to a number of reasons, including slight shifts in the position of the array (e.g., when the animal moved around in its home cage) or a build up of glial tissue over time around the implanted electrodes. To retain the largest number of neurons possible, we made the assumption in our modeling framework that each session had a new set of neurons (Ext. Data Fig. 1). The number of neurons across recording sessions and animals ranged from 21 to 89 neurons with a median of 58 neurons; exact numbers are in Extended Data Table 1. Similar to previous studies [12, 40], we took spike counts in 100 ms time bins starting 50 ms after stimulus onset to account for synaptic delays to V4.

#### Active fixation task

To record responses to a large number of images, we trained each animal to perform a simple active fixation task. Each trial began with the presentation of a central blue fixation dot. The animal held an initial fixation for 150 ms. Following this, a sequence of 6 or 8 images (depending on the animal) was presented while the animal maintained fixation; images overlapped with the receptive fields of the V4 neurons. The side length of each square image was 11.2°, 10.6°, and 8.0° for monkeys WE, PE, and RA. The images were positioned to be 12.7°/5.0°, 4.4°/6.2°, and 3.8°/3.8° below/to the right of central fixation dot for monkeys WE, PE, and RA. Each image was displayed for 100 ms with a 100 ms gray screen between images. Following this image sequence, the central fixation dot vanished and a target dot appeared∼ 10° from the location of central fixation. To receive a liquid reward (i.e., a ‘correct’ trial), the animal had to make a saccade to the target. Breaking fixation or failing to make a saccade to the target dot resulted in no reward and a 1 s time out before the next trial began. The gaze of the animal was tracked using an infrared eye tracking system (EyeLink 1000; SR Research, Ottawa, Ontario) and monitored online by experimental control software to ensure fixation and to report the animal’s choice target. The display was a gamma-corrected flat-screen cathode ray tube monitor positioned 57 cm from the animal’s eyes with a resolution of 1024*×* 768 pixels, refreshed at a frame rate of 100 Hz. The background of the display was 50% luminance (gray). All animals performed the task well with many thousands of flashed images in each session.

Before each session, the set of images to show were chosen either randomly from a large data set, in a closed-loop manner, or by synthesizing images via the model (see ‘Visual stimuli’ section). The total number of unique images per session, ranging from 400 to 3,000 images and typically ∼ 2,000 images, was chosen to ensure enough repeats per image. All images were shown once in randomized order; for subsequent repeats, we showed the sequence of all images again but with a different randomized order. Each session lasted for multiple hours and ended when the animal showed signs of satiation. The number of repeats per image for each session ranged from 2 to 24 repeats; exact values are listed in Extended Data Table 1. To increase the number of repeats, we included any repeat for which the animal remained fixating on the central dot while the image was displayed; this included trials in which the animal may have broken fixation early, ending the trial and receiving no reward (i.e., an error trial). We discarded responses to images with only one repeat. The total number of sessions was 50 (7, 29, 14 sessions for monkeys WE, PE, RA). Of these sessions, 44 sessions were used to train our deep ensemble model; 1 session of entirely normal images was used to validate any model hyperparameters; 4 sessions of entirely normal images were used to test model predictions (1 session for monkey WE, 1 session for monkey RA, and 2 sessions for monkey PE recorded 4 months apart); and 1 session of images from a previous study [40] was used for comparison (Ext. Data Fig. 2). Images were unique across the validation and test sessions, and no model had access to any of these images during training (i.e., held out images).

### Visual stimuli

Our experiments involved many different types of images. These types included normal images, images chosen by active learning, gaudy images, probe images used for causal testing, synthesized images (generated with DNNs), and artificial images (e.g., sinusoidal gratings, dot images, etc.). All images were 112 *×* 112 pixels with RGB color channels (i.e., the images were not restricted to grayscale). The types of images (and number of each type) shown for each session can be found in Extended Data Table 1—the total number of images per image type varied. We briefly describe each type of image here; examples of images are shown in Extended Data Fig 7.

1. *Normal images*: We define normal images as those coming from our dataset of 12 million “natural” colorful images subsampled from the Yahoo Flickr Creative Commons 100 Million Dataset (YFCC100M) [107], which contains ∼100 million images uploaded by users to Flickr between 2004 and 2014. Images are unlabeled (i.e., no content information) and need not contain an object. After sampling an RGB image from YFCC100M, we first randomly cropped the image to have an equal number of row and column pixels and then downsized the image to 112 × 112 pixels. We chose YFCC100M primarily to ensure that our images used for training and testing were different from those in ImageNet [108], as the task-driven DNNs that we chose for features (e.g., ResNet50) were trained on ImageNet images.
2. *Images chosen adaptively by active learning*: In some of the sessions, a portion of the images were chosen adaptively via a closed-loop method called active learning (also known as optimal experimental design) [43, 44, 45, 42, 54, 109]. To choose these images, we trained our deep ensemble model (see below) on all past recording sessions. Then, for each of 500,000 candidate images randomly chosen from our image data set, we computed the disagreement across ensemble members as the variance of predicted responses, averaged over neurons from the last recording session. We chose the images with the largest ensemble disagreement to show in the next session.
3. *Gaudy images*: In our previous work, we identified a class of images that were adept at efficiently training DNN models of visual cortex, even better (in simulations) than images chosen with active learning [45]. We called these images “gaudy” images for their high-contrast edges and over-the-top colors. To generate a gaudy image, we first randomly chose an image from our image data set. Then, for each pixel intensity, if that pixel intensity was greater than the average intensity across pixels for the same RGB channel, we set its value to 255; else 0. Thus, each pixel was one of 2^3^ different colors (black, white, red, green, blue, etc.).
4. *Probe images used for causal testing*: To test our compact model predictions, we probed each compact model in three different ways to identify its preferred stimuli. First, we identified the image from 500,000 candidate normal images in our image data set that maximized the output of the compact model (Fig. 2**b**). Second, we synthesized the image via gradient methods that maximized the output of the compact model (Fig. 2**c**; see ‘Maximizing synthesized images’ section). Third, we synthesized adversarial images that either maximized or minimized the output response of the compact model (Fig. 2**d**-**f**; see ‘Causal tests’ section).
5. *Maximizing normal and synthesized images of deep ensemble model*: We identified maximizing normal images and maximizing synthesized images for the deep ensemble model in the same way we identified the probe images for the compact models.
6. *Artificial images*: In a small number of sessions, we presented artificial stimuli. For one session, we presented static sinusoidal gratings varying in orientation (from 0° to 162°, 14 equally-spaced orientations in total) and spatial frequency (from 1/50 pixels/° to 1/2 pixels/°, 14 equally-spaced frequencies in total). In another session, we presented “dot” images which varied in dot location, dot size, and number of dots (Ext. Data Fig. 6).

### Predicting V4 responses

Given a set of model features taken either from a task-driven DNN or our deep ensemble model (described in later sections), we estimated the extent to which the features predicted V4 responses via a linear mapping. We follow recent work that uses a factorized linear mapping [40, 110], taking advantage of spatial and filter information. Specifically, consider the output features (or activity maps) of a convolutional layer for one image as a tensor *X* of shape *p × p × K* for *p* pixels and *K* convolutional filters. Fitting the weights of a linear regression to one V4 neuron would result in weight tensor *W* of shape *p × p × K*. With a factorized linear mapping, we first spatially readout each activity map with the same spatial filter (*W*_spatial_ with shape *p × p*) and then linearly combine these readouts across filters (*W*_mixing_ with shape *K ×* 1) via the following equation:

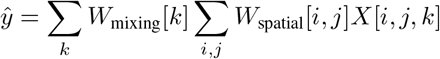

This factorized linear mapping reduces the number of weights to *p × p* + *K*; in other words, *W*_spatial_ and *W*_mixing_ together form a low-rank approximation of *W*, the weight tensor from standard linear regression. We regularized the weights of the factorized linear mapping with an L2 penalty; the hyperparameter *α* factor was fit with the validation session. We confirmed that our fitting procedure of the factorized linear mapping reproduces previously-reported prediction performance on a publicly-released data set of V4 responses [40] (see Ext. Data Fig 2).

We held out 4 recording sessions for which we presented ∼ 1,000 colorful, natural images per session (images were unique across sessions) with ~10 repeats per image (see Ext. Data Table 1). For each session, we used 4-fold cross-validation; we trained the factorized linear mapping on 3/4 of the heldout image-response pairs and computed the predicted responses for the remaining 1/4 images. We then collected the predicted responses for all cross-validation folds as one vector 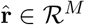or all *M* images of the session.

To assess prediction performance, standard practice is to account for repeat-to-repeat variability by normalizing the explained variance between the model and repeat-averaged responses 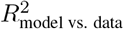 by the explained variance between two sets of repeat-averaged responses where repeats are split into two groups 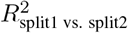 [12, 30]. The 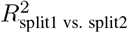 can be thought of as a noise ceiling for the largest possible value of 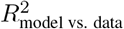 . Recent work has shown that this estimation procedure is biased, leading to overly-optimistic estimates, and proposes an unbiased, noise-corrected 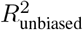 [106], which we use in this work (see Ext. Data Fig. 2 for comparisons). For a neuron’s repeat-averaged response **r**_*i*_ and the model’s predicted response 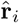 to the *i*th image, the 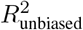 is computed as follows:

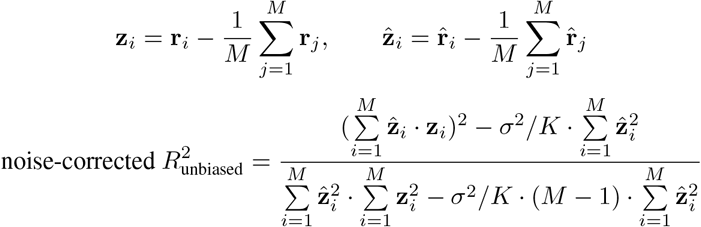

where *M* is the number of images and *K* is the number of repeats per image. The left terms in both the numerator and denominator are the same as found in computing standard *R*^2^, and the right terms in both the numerator and denominator subtract off the bias caused by repeat-to-repeat variability *σ*^2^. Because different images may have a different number of repeats (depending on how many trials were completed by the animal), we took the unbiased estimate of *σ*^2^ for each image and averaged across images: 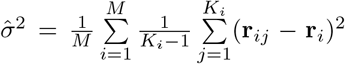 where K_*i*_ is the number of repeats for the *i*th image and **r**_*ij*_ is the response for the *i*th image and *j*th repeat. In addition, we set *K* to the average number of repeats across images.

We computed 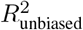 between the predicted responses 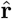 and recorded responses **r** across all images in the session (i.e., we did not compute 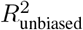 per fold and average across folds), as we found this better avoided the effects of outliers with small numbers of images (*<* 500 test images per fold). When directly compared, we found that estimates for the previously-used 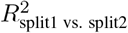 were on average ∼0.15 larger than estimates for 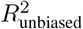 (Ext. Data Fig. 2), suggesting that current models in the field are not as predictive as once thought. We call 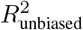 as noise-corrected *R*^2^ in the rest of the paper.

### Task-driven DNN models

We predicted held out V4 responses with task-driven DNN models (Fig. 1**b**, trained to perform object recognition on ImageNet [108]), as previous studies have found the internal layers of these task-drive DNNs to be predictive of V4 responses [1, 40]. We chose all DNNs readily available to download in Keras [111]; each DNN differed in architecture, optimization, and classification performance on ImageNet. To identify the layer used to predict V4 responses for each DNN, we computed the cross-validated noise-corrected *R*^2^ between each layer’s activity and V4 responses from the validation session using ridge regression and chose the layer—typically one of the middle layers, consistent with previous studies [12, 30]—with the largest *R*^2^ (see ‘Predicting V4 responses’ section). The DNNs and identified layers (using the names as defined in Keras) were as follows: VGG19, block4_pool [112]; InceptionV3, mixed4 [113]; InceptionResNetV2, mixed_6a [114]; DenseNet169, `pool3_pool` [115]; ResNet50, `activation_33` [66]; MobileNet, `conv_pw_9_relu` [116]; NASNetMobile, `activation_104` [117]; NASNet-Large, `activation_128` [117]; Xception, `add_7` [118]. Input images were resized and processed according to each DNN’s data processing procedure. We found the average prediction performance of noise-corrected *R*^2^ *≈* 0.4 (Fig. 1**b** and Ext. Data Fig. 2) comparable to that reported in previous studies [12, 38, 30]. An untrained task-driven model (ResNet50 with randomly-initialized weights) had a much worse noise-corrected *R*^2^ = 0.13, suggesting that training on a relevant task (in this case, object recognition) is needed for good prediction of higher-order visual cortical neurons.

### Deep ensemble model

Our deep ensemble model (Fig. 1**a**) relied on transfer learning and ensemble learning. To compute the predicted response for an image, the image was first processed and passed through ResNet50 until layer `activation_33` which outputs a 14 *×* 14 *×* 1, 024 tensor of activity maps (each with shape 14 *×* 14) for 1, 024 channels. This feature tensor was passed as input into an ensemble of small DNNs; each ensemble DNN received the same feature tensor as input. Each ensemble DNN outputs a vector of predicted responses for *n* neurons; these vectors were averaged across the ensemble to get the final predicted response vector.

#### Architecture of ensemble DNN

We briefly describe the architecture of the ensemble DNN; Extended Data Figure 1 contains a detailed diagram. The input is a 14 *×* 14 *×* 1, 024 feature tensor from a middle layer of ResNet50. The ensemble DNN’s first layer reduces the number of input channels from 1,024 to 512 by taking linear combinations of the input channels (i.e., a kernel shape of 1 *×* 1); this is followed by batchnorm [119] and ReLU activation operations. The second layer is a separable 2-d convolution [116] of 512 filters with kernel shape 3*×* 3 and a stride of 2. The output of layer 2 is a 7 *×* 7 *×* 512 tensor. Layers 3 through 6 have the same following skip connections, inspired by ResNet [66]. This comprises 1) a separable 2-d convolution (256 filters with kernel shape 3 *×* 3) followed by batchnorm and ReLUs, 2) a separable 2-d convolution (256 filters with kernel shape 3 *×* 3) followed by batchnorm and ReLUs, and 3) a convolutional layer to expand the dimensionality (512 filters with kernel shape 1 × 1). The output, a tensor of shape 7 *×* 7 *×* 512, is added to the layer’s input (i.e., a skip connection) and is used as input to the next layer. The final output embedding (i.e., the output of layer 6) is a tensor with shape 7 *×* 7*×* 512. This embedding is passed through a factorized linear mapping (with a mixing stage and a spatial pool stage, see ‘Predicting V4 responses’) to get predicted V4 responses. We chose this architecture and hyperparameters after a large architecture search (e.g., Fig. 1), validating with one held out validation recording session (different from the 4 recording sessions used for testing performance).

#### Training

We trained the deep ensemble model on 45 recording sessions. Each recording session had its own factorized linear mappings trained independently from any other session; all other model weights were shared across sessions (see Ext. Data Fig. 1). Responses for each V4 neuron were repeat-averaged and then z-scored to balance gradients when predicting multiple V4 neurons. We optimized the model using SGD with learning rate *η* = 20, momentum *ν* = 0.7, learning rate decay *δ* = 0.85 and a batch size of 64; optimizing with Adam [120] did not perform well when training across sessions. We took the model across training epochs with the best prediction performance evaluated on a validation recording session (i.e., early stopping).

#### Boosts in performance versus task-driven models

Our deep ensemble model was highly predictive of V4 responses to held out images with a median noise-corrected *R*^2^ = 0.60 (median *R*^2^ = 0.55, 0.63, 0.63, 0.58 for the 4 held out recording sessions from monkeys WE, PE, PE, and RA, respectively). We performed 4 analyses to assess which design decisions in our modeling framework led to such a large increase in prediction performance (Fig. 1**c**-**f**). For the first analysis, we asked whether allowing for nonlinearities between ResNet50 features and V4 responses led to better prediction. We trained a purely linear mapping that comprised the ensemble model’s first 2 layers with no ReLUs (Fig. 1**c**, ‘linear’), as well as ensemble models with different numbers of skip connection layers (Fig. 1**c**, ‘num. ReLU layers’). We chose 4 skip connection layers for our final model. For the second analysis, we varied the number of training sessions *k* by randomly choosing a subset of *k* sessions out of the 45 sessions used for training (Fig. 1**d**). For the third analysis, we varied the number of ensemble DNNs by training a model with 40 ensemble DNNs and taking subsets of ensemble DNNs; for each subset, we recomputed prediction performance (Fig. 1**e**). We chose 25 ensemble DNNs for our final model to ensure good prediction and estimation of ensemble disagreement (used for active learning). For the fourth analysis, we assessed to what extent training the ensemble model on images chosen by active learning in closed-loop experiments and gaudy images (see ‘Visual stimuli’ section) better predicted V4 responses than training on randomly-chosen normal images (Fig. 1**f**). We only considered training sessions in which both types of images were shown (30 sessions total). We equalized the number of images for both types by random sampling and re-trained the deep ensemble model on each type separately (using a learning rate *η* = 1.5 to account for the smaller amount of training data). We varied the number of training sessions by taking random subsets of sessions, training a new ensemble model for each subset. Importantly, we tested prediction performance on the same 4 held out sessions as for the other analyses; these sessions only had responses to randomly-chosen normal images. In other words, training on active learning/gaudy images while testing on normal images represents an out-of-distribution prediction problem; only if these active learning/gaudy images better emphasize important features of normal images would training on them achieve better prediction performance than that of training on randomly-chosen normal images [45].

### Identifying compact models

Ideally, we would start with a compact DNN model and directly fit its weights to the real data. However, when the amount of training data is limited (as in our case), this often leads to overfitting—indeed, training a small DNN led to poor performance (noise-corrected *R*^2^ = 0.11). A large DNN model, trained on the same data, overcomes this overfitting by having access to many subnetworks or “lottery tickets” [34]. Via pruning techniques [36, 46, 47, 48, 49], one can identify a small subnetwork (or winning ticket) that retains the same prediction performance as the large model. Motivated by this approach, we decided to take our highly-predictive deep ensemble model and find a compact model with as few parameters as possible yet the same prediction performance as that of the deep ensemble model.

To identify a compact model (one for each heldout V4 neuron), we first obtained the predicted responses from the deep ensemble model to 12 million images from our image data set. For each heldout V4 neuron, we trained a factorized linear mapping (see ‘Predicting V4 responses’ section) from the deep ensemble model’s output embeddings to V4 responses using half of the image-response pairs for that neuron’s session—the other half was held out to measure prediction performance (noise-corrected *R*^2^s in Fig. 1**h**). We then performed two model compression techniques used in deep learning: knowledge distillation and pruning. Knowledge distillation trains a single DNN to predict the averaged output of an ensemble of DNNs [35]; the single DNN overcomes overfitting because unlike the relatively small number of image-V4 response pairs ( ∼ 2,000 per session), we train the distilled DNN on ensemble responses to 12 million images. Pruning takes a trained DNN and ablates unnecessary weights (i.e., set their values to 0) [36, 46, 47, 48, 49]; DNNs with up to 90% of their weights pruned have been found to be as predictive as the original unpruned DNN [34]. We employed both of these techniques to identify compact models.

We first performed distillation on our ensemble model. We initialized a ‘distilled’ DNN model with the following architecture. The (re-centered) input RGB image has shape 112 *×* 112 *×* 3. The first layer is 2-d convolutional comprising 100 filters with kernel size 5 *×* 5 followed by batchnorm and ReLUs. Layers 2-5 are each 2-d separable convolutions comprising 100 filters with kernel size 5 *×* 5 followed by batchnorm and ReLUs; layers 2 and 3 take a stride of 2. The final dense ‘spatial readout layer’ linearly maps the output activity of layer 5 (of shape 28 *×*28 *×* 100) to the scalar V4 response. We chose all layers to have 100 filters after a hyperparameter sweep. We trained the distilled model on one pass of 12 million image-response pairs, where responses were the predicted responses from our ensemble model, using Adam [120] to minimize the mean squared error with learning rate *η* = 10^*−*4^. For every 500,000 training images, we smoothed the kernel weights of layers 2-5 and the spatial readout layer with a gaussian filter (*σ* = 0.5 pixels) for better interpretability of the kernel weights.

The distilled models were substantially smaller than our deep ensemble model (600 filters versus ∼ 150,000). However, we wondered whether different V4 neurons could be explained by compact models with different numbers of filters. For example, an edge-detecting V4 neuron may need fewer filters than a curve-detecting V4 neuron. To test this, we developed a channel-wise pruning technique that ablated entire filters (as ablating individual weights could possibly lead to many filters with sparse weights, making the non-smooth kernel weights difficult to interpret). We developed and performed the following pruning technique, inspired by hard filter pruning from previous studies [48, 49]. We first pass 5,000 randomly-chosen normal images as input and collect the output activity for each convolutional layer and the spatial readout layer. Then, for each layer, we order the filters based on the variance of each filter’s activity summed over spatial locations. We then identify the largest subset of lowest-variance filters that, once ablated, still explains at least 90% of the variance of the original output activity for that layer. In other words, we remove filters with either weak activity or whose activity is unlikely to reach downstream layers. We perform this procedure first for the spatial readout layer (i.e., the deepest layer), followed by the preceding layer until layer 1 is reached; reversing this layer order (i.e., pruning early to late layers) leads to pruned models with ∼ 100 extra filters (Ext. Data Fig. 2). We then re-train this pruned network in the same way as the distilled network. We call the re-trained pruned network a ‘compact’ model, one for each held out V4 neuron (219 in total). We note that the number of convolutional filters in layers 1 and 2 must be equal by definition as consequence of having a convolutional layer followed by a separable convolutional layer (Fig. 3**a** and Ext. DataFig. 1).

### Maximizing synthesized images

To identify the visual features that a compact model prefers, one approach is to search for the image that maximizes the output response of the model. A simple search involves computing the model’s output response for a large pool of candidate images and then choosing the image that evokes the largest output [50]. However, because the pool of candidate images is finite, not all possible images are considered. A more expressive search is to synthesize the maximizing image via gradient ascent [40, 51]. This takes advantage of the fact that the mapping of the compact model is entirely differentiable. The search begins with a white noise image where each pixel intensity’s value is drawn from a Gaussian with mean 128 and standard deviation 50 (clipped between 0 and 255). This initial image is passed through the model to compute its output. We then take the gradient of the output with respect to the input image and pass this gradient back through the network (i.e., backpropagation), updating the image with the gradient ascent rule: **x**_next_ = **x**_current_ + *η▽***f**(**x**) for image **x**, gradient of model output ▽**f**(**x**), and step size *η* (in our analyses, *η* = 10). We perform this for 1,000 steps or until the maximized output did not change. Because moving along the gradient direction may lead to a synthesized image far outside the natural image manifold, we smooth the gradient at each step (Gaussian filter with *σ* = 1) and smooth the image (Gaussian filter with *σ* = 1) every 50 epochs, similar to previous approaches [40, 51].

### Causal testing of the compact models

We made the following three predictions of our compact models that we causally tested in follow-up experiments. We ran these experiments in two monkeys (PE and RA). We trained the deep ensemble model on any previously-recorded sessions from any animal. We then identified a compact model for each V4 neuron on a heldout session in which only normal images were shown, following the same process as described in “Identifying compact models”. We obtained three different predictions from the compact models. First, we identified the top maximizing normal image from 500,000 normal images for each compact model (Fig. 2**b**; 1 image for monkey PE and 10 images for monkey RA per compact model). Second, we identified the top maximizing synthesized image following the same process as outlined in “Maximizing synthesized images” (Fig. 2**c**; 1 image for monkey PE and 10 images for monkey RA per compact model; each image began with a different white noise image). Third, we identified adversarial images [63] for each compact model (Fig. 2**d** and **f**; 12 images for monkey PE and 10 images for monkey RA per compact model). We computed the adversarial images that either maximized the model’s output response (via gradient ascent) or minimized the model’s output response (via gradient descent). For each adversarial image, we continued the optimization procedure until either 50 gradient updates were completed or the average intensity difference (absolute valued) between pixels of the initial image and the synthesized adversarial image passed a threshold pixel intensity of 10 (i.e., an average change of 0.04 for normalized pixel intensities that range from 0 to 1). We set the intensity threshold higher than intensity differences typically observed in deep learning studies [61, 63] to ensure the animal could make a perceptual difference between the two images; we were not interested in identifying the smallest image perturbations that led to changes in V4 responses (for IT neurons, see [64]) but rather if the compact models could be used to slightly perturb an image (out of an inifinite number of possible perturbations) to elicit large changes in V4 responses.

We presented these probe images in succeeding sessions (one session for both maximizing normal and synthesized images and one session for adversarial images). For comparison, we also presented a number of normal images within each session (250 images for monkey PE and 660 images for monkey RA). After recording V4 responses, we assessed if the probe images did elicit changes in responses compared to normal images. Because there was no guarantee the recorded neurons on succeeding sessions would be the same as those targeted by the compact models, we first had to match each V4 neuron with a compact model. To do this, for every V4 neuron, we computed the noise-corrected *R*^2^ between that neuron’s responses and the compact model’s predicted responses on all presented images except the maximizing/adversarial images. The compact model that matched to a V4 neuron was the one with the largest *R*^2^. We then computed the repeat-averaged responses of a V4 neuron to the probe images for the matched compact model (Fig. 2**b**-**f**).

### Similarity of filter kernels

A coarse measure of whether two different compact models have similar computations in the same layer is to assess whether the two models have similar convolutional filter kernels. We devised a kernel similarity metric where a value close to 1 indicates that each convolutional filter in one model has a corresponding convolutional filter that matches in its pattern of kernel weights (i.e., the 5*×* 5 weight matrix for each convolutional filter). A kernel similarity close to 0 indicates that no matching filters are present.

Ideally, we would like to compare kernel similarity across layers to make statements such as layer *i* has more similar filters than those of layer *j*). To do this, we must account for the different number of filters across layers (Fig. 3**a**). We devised a kernel similarity metric to account for this difference, defined as follows. For a given layer, we randomly choose two non-overlapping sets of 100 filters each. For each pair of filters between the two sets, we compute the dot product of their kernel weights (each normalized to unit norm) and take its absolute value. Then, in an iterative manner, we select the filter pair with the largest value and select the next filter pair such that a filter can only be chosen for a single pair. After no pairs remain, we define the kernel similarity as the average value across all selected pairs. We perform this procedure for 500 runs; the mean and s.d. are reported in Figure 3**b**.

Because the kernel size of layer 1 filters was 5 *×* 5 *×* 3 to account for the 3 RGB channels, we averaged each filter’s kernel weights over the RGB channels, resulting in a 5 *×* 5 filter. Still, it was not fair to compare kernel similarities between layer 1 and any other layer, as layer 1 was not smoothed during training (whereas layers 2-5 were smoothed), leading to more “jagged” kernel weight patterns. In addition, we could not compare kernel similarities between the spatial readout layer and any other layer, as the filters in the spatial readout layer were 28 *×* 28, much larger than the 5 *×* 5 filters in the other layers. For these reasons, we only compare kernel similarities among layers 2-5. For reference, we computed the kernel similarity if all filters had kernel weights drawn from a standard normal.

### CKA similarity

To compare the internal representations between two compact models (Fig. 3**c**), we employed Centered Kernel Alignment (CKA) similarity, a commonly-used metric to compare representations between two deep neural networks [71]. CKA similarity is computed as follows. Consider the response activity of two representations 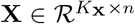 and 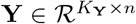 for the same *n* images, where *K*_**X**_ and *K*_**Y**_ are the number of feature variables in each representation (*K*_**X**_ and *K*_**Y**_ need not be equal). We assume that **X** and **Y** are centered—the means of each row are 0. We then define CKA similarity (using a linear kernel as in [71]) as follows:

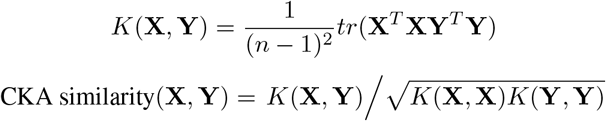

where *tr* denotes the trace operation. A CKA similarity close to 1 indicates two representations are highly similar; a value close to 0 indicates the representations are different.

CKA similarity has several nice properties, including invariance to orthogonal linear transformations (e.g., if the rows of **X** were a permutation of the rows of **Y**) and invariance to isotropic scaling (e.g., if **X** = *c***Y** for some scalar *c*). As opposed to a similarity metric based on canonical correlation analysis [121], CKA similarity will be larger if the eigenspectra of the two representations are similar—in other words, CKA similarity captures if two representations have the same dominant dimensions or not. In addition, when both representations have one variable each, CKA similarity is equivalent to the squared Pearson correlation (i.e., the signal correlation squared between the outputs of two models). Other similarity metrics exist [121, 122]; however, because almost all of these similarity metrics rely on the same principles of invariance to linear transformations and isotropic scaling, we do not suspect our conclusions to differ using a different similarity metric. We computed the CKA similarity between all pairs of models for each of the 5 layers, as well as the model’s output, in response to 10,000 randomly-chosen normal images (Fig. 3**c**).

### Compact model shared across all V4 neurons

Given our finding that the early layers of the compact models had similar convolutional filters (Fig. 3**b**) and similar representations across V4 neurons (Fig. 3**c**), we reasoned that we could identify a ‘shared’ compact model that predicted all 219 heldout V4 neurons together. Specifically, we wondered how many filters *k* were needed in the first three layers of this shared compact model. The architecture of the shared compact model resembled that of a compact model before pruning (i.e., the ‘distilled’ model, see Ext. Data Fig. 1). Namely, layer 1 was a 2-d convolutional layer comprising *k* filters with kernel size 5 *×* 5 *×* 3 (for 5 *×* 5 image patches and three RGB channels) followed by batchnorm and ReLUs. Layers 2 and 3 were 2-d separable convolutional layers comprising *k* filters with kernel size 5 *×* 5 and a stride of 2 followed by batchnorm and ReLUs. Layers 4 and 5 were the same as layers 2 and 3 except always with 100 filters. The output embedding for *m* images is a tensor with shape *m ×* 28 *×* 28 *×* 100. We map this embedding to the responses of *n* V4 neurons via a factorized linear mapping (see ‘Predicting V4 responses’). This architecture allowed us to vary the number of filters in the first three layers *k* while allowing specialization for layers 4 and 5. Similar to how we trained each compact model, we trained the shared compact model on predicted responses of our deep ensemble model to 12 million images; to minimize our loss function of mean squared error, we used the Adam optimizer [120] with a learning rate of *η* = 10^*−*4^. For every 500,000 training images, we smoothed the kernel weights of the convolutional filters in layers 2-5 and the 28 *×* 28 kernel weights in the spatial pooling stage of the factorized linear mapping (Gaussian smoothing with smoothing factor *σ* = 0.5 pixels). We computed the noise-corrected *R*^2^ for each heldout V4 neuron on the heldout half-split of images (the other half of images was used to linearly map the deep ensemble model to V4 responses). We varied the number of early layer filters *k* (Fig. 3**e**) and trained a new shared compact model for each *k*. The noise-corrected *R*^2^ values reported for this analysis slightly differ from those in Fig. 1**b** because here we held out half of a V4 neuron’s responses to compute prediction performance while in Fig. 1**b** we perform cross-validation for the responses. For reference, we fixed the randomly initialized kernel weights for the *k* convolutional filters of layers 1-3, trained all remaining weights of the shared compact model (including the ‘mixing’ weights of the separable convolutions), and re-computed prediction performance (Fig. 3**e**, ‘untrained shared filters’). If prediction performance of a shared compact model with untrained shared filters matched that of a shared compact model with trained shared filters, it suggests that the similarities among early layers does not arise from shared kernel weights. We found this was not the case (Fig. 3**e**, ‘trained shared filters’ line above ‘untrained shared filters’ line). However, as expected, increasing the number of filters *k* led to a smaller difference between prediction performances because the untrained, random kernel weights form an overcomplete basis of filters (i.e., with enough random linear projections, all information about the input is retained); for a large enough *k*, both models (with trained and untrained convolutional filters) would achieve similar prediction performance.

### Dot detector characterization

To identify a compact model that exemplified a “dot detector”, we assessed the preferred stimuli for each compact model (e.g., the maximizing normal and synthesized images) and chose the one that most resembled a dot detector. To ensure the chosen compact model was a dot detector, we devised an artificial stimulus set in which we varied the dot location, size, and number (Fig. 4**b** and Ext. Data Fig. 5). For dot location, we varied the location of a dot with a small size (5 pixel radius) for 28 *×* 28 equally-spaced locations; the model’s preferred dot location is the dot location with the largest response. For dot size, we placed a dot at the model’s preferred location and varied its radius from 2 pixels to 22 pixels. For dot number, we placed *k* dots randomly within 25 pixels of the preferred location and varied *k* from 1 to 10 dots. For each dot number, we generated 10 instances and took the mean response of the model.

To identify which filters contributed to the chosen model’s selectivity to dot size, we considered the model’s output responses **r**_no ablation_ to the dot size stimuli versus the model’s output **r**_*¬*L*ℓ*F*j*_ when we ablated the *j*th filter from layer *ℓ* (i.e., set its kernel weights to 0). If a filter contributed to dot size selectivity, we would expect to see a large change in the model’s output. To quantify this, we defined a dot size invariance (DSI) metric for the *j*th filter of layer *ℓ* as follows:

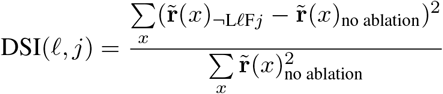

where *x* denotes which dot size stimulus, and 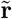 denotes the residual model response after the mean model response across all dot size stimuli was subtracted. We found this re-centering to be necessary, as ablating some filters produced an additive offset change in response but otherwise left the dot size selectivity of the model intact. To identify which filters in layer 3 were necessary (Fig. 4**e**), we cumulatively ablated layer 3 filters by the following greedy procedure. First, we identified the individual filter that, once ablated, yielded the *smallest* increase in DSI (i.e., the filter that *least* contributed to the model’s dot selectivity). We repeated this process for the remaining unablated filters while keeping all chosen filters ablated. At some point, DSI must increase, as ablating all filters leads to DSI = 1. Thus, the filters chosen last in this process (i.e., the ones where DSI starts to increase) contribute most to the model’s dot size selectivity. We investigated the 12 filters to be chosen last (rightmost dots in Fig. 4**e**), from which we found 6 filters that carried out a simple computation (Fig. 4**f**-**h**); the remaining 6 filters appeared to be redundant or had weaker effects.

We also found that the dot-detecing compact model preferred multiple dots with varying positions (Fig. 4**a**). In a similar analysis, we identified the circuit mechanisms contributing most to dot number selectivity (Ext. Data Fig. 5).

### Statistical analysis

Significance testing was performed with two-tailed permutation tests unless otherwise states. If the actual statistic was greater or less than all “permuted” statistics under the null hypothesis, the *p*-value was set to 0.002, equal to 1*/n* where the number of runs *n* = 500. Bootstrapped confidence intervals of medians were computed by 500 runs of re-estimating the median by sampling from the data with replacement. Numbers of images, repeats, neurons, and sessions are listed in Extended Data Table 1.

## Extended Data Figures

**Extended Data Figure 1:**
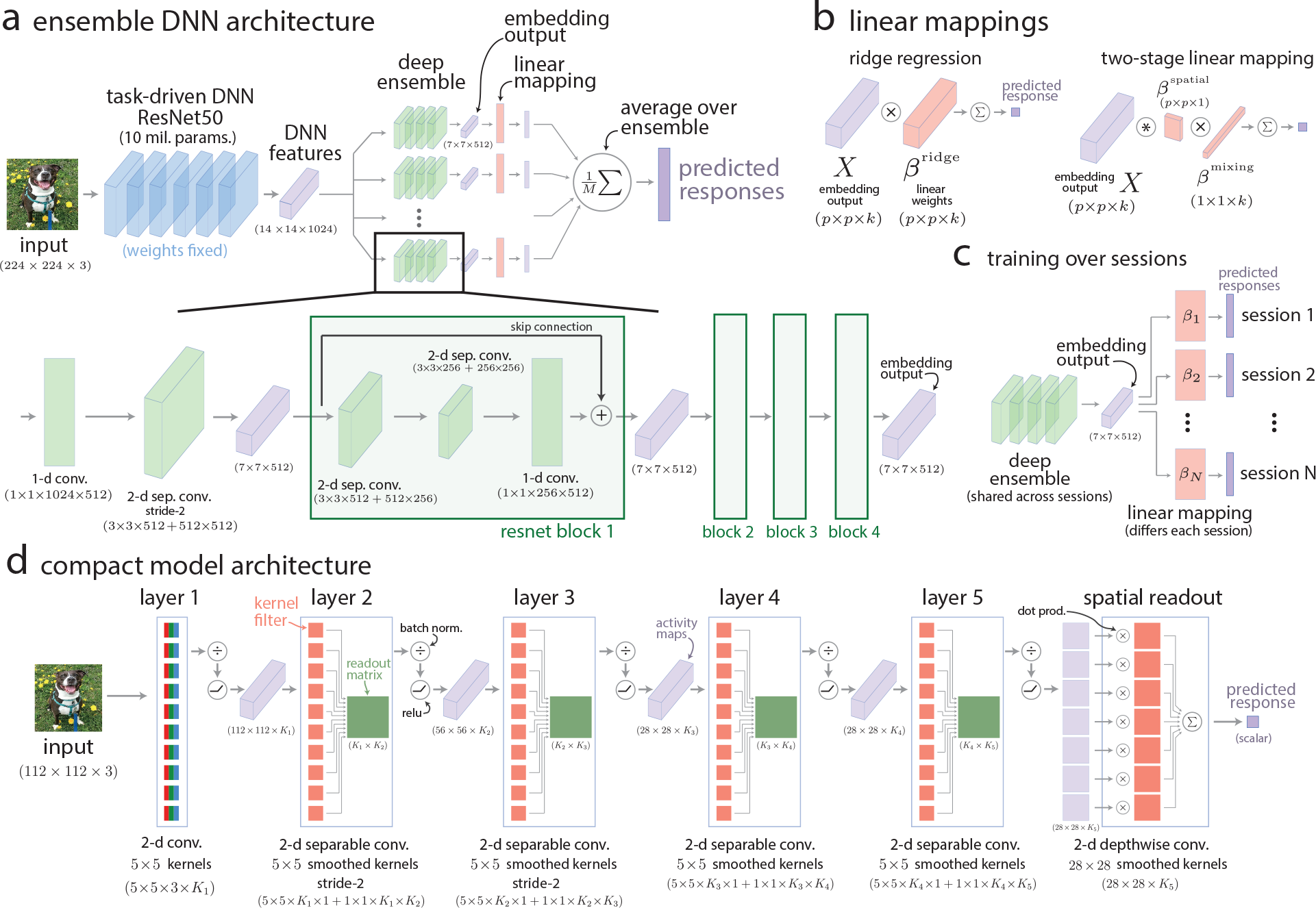
Detailed diagrams of deep ensemble model and compact model network architectures. **a**. Detailed diagram of our deep ensemble model, which uses the early layers of a task-driven ResNet50 and an ensemble of ResNet-like DNNs. Each ensemble DNN has skip-connection blocks and relies on separable convolutions to reduce the number of parameters. The shapes of each output activity tensor as well as the shapes of the weight tensors are in parentheses nearby their corresponding layer. See Methods for details. **b**. Diagrams of linear mappings between a model’s embedding output activity *X* (of shape *p × p × k* for *p* downsampled pixels and *k* channels/filters) and V4 responses. Ridge regression fits a weight matrix *β*^ridge^ of shape *p × p × k* with *L*_2_ regularization (top). The predicted response 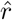 is computed from embedding *X* as the following:

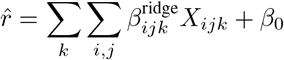

where *β*_0_ is an offset term. The factorized linear mapping [110] fits two weight matrices: a mixing matrix *β*^mixing^ of shape *k ×* 1 that integrates or “mixes” information across filters and a spatial pooling matrix *β*_spatial_ of shape *p × p* that integrates spatial information. The predicted response 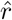 is computed as the following:

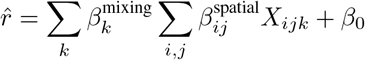

This factorized mapping can be thought of as a low-rank approximation to ridge regression. Comparing the number of parameters between the two mappings reveals that the number of parameters of ridge regression (*p p k*) is substantially greater than that of the factorized linear mapping (*p · p* + *k*). Given that the linear mapping needs to be fit to a small amount of training data (typically *<* 1, 500 images), the factorized linear mapping will less likely overfit due to its smaller number of parameters (Ext. Data Fig. 2). **c**. Although the recording electrode array was chronically implanted, we could not be entirely certain that we recorded the same V4 neuron across two or more sessions. Thus, we assumed that each recording session was a new sampling of V4 neurons (with the caveat that some neurons were likely present in multiple sessions). To build this assumption into the model, we gave each recording session its own linear mapping between the embedding output of the deep ensemble model and that session’s V4 responses (*β*_1_, *β*_2_, …, *β*_*N*_ for *N* sessions). When training the deep ensemble model on the *i*th session, we set the weights of *β* to 0 (but kept all other weights of the deep ensemble the same) and performed stochastic gradient descent end-to-end; we found that resetting *β* with previously-trained *β*’s led to overfitting and worse performance. To evaluate the model’s predictions on a held out session, we trained *β* on a portion of the image/response pairs and predicted the remaining pairs in a cross-validated manner. **d.** Detailed diagram of a compact model, including the separable convolutional layers, batchnorms, and ReLUs. The *i*th layer has its own number of filters *K*_*i*_ except for layers 1 and 2 which have the same number of filters (*K*_1_ = *K*_2_) due to layer 1 being fully convolutional. We illustrate separable convolutions in two parts: the convolutional filters (red squares) and the mixing weights (green matrices). Each convolutional filter processes the activity map of one input filter, making the number of convolutional filters in the *ℓ*th layer equal to *K*_*ℓ−*1_, the number of output channels for the (*ℓ −* 1)th layer. Each mixing matrix linearly combines the output of these convolutional filters across filters and takes shape *K*_*ℓ−*1_ *× K*_*ℓ*_. Layer 1 is a fully convolutional layer, and the spatial readout layer is a dense layer but can be thought of as a set of spatial receptive fields whose output is summed together. See Methods for further details.

**Extended Data Figure 2:**
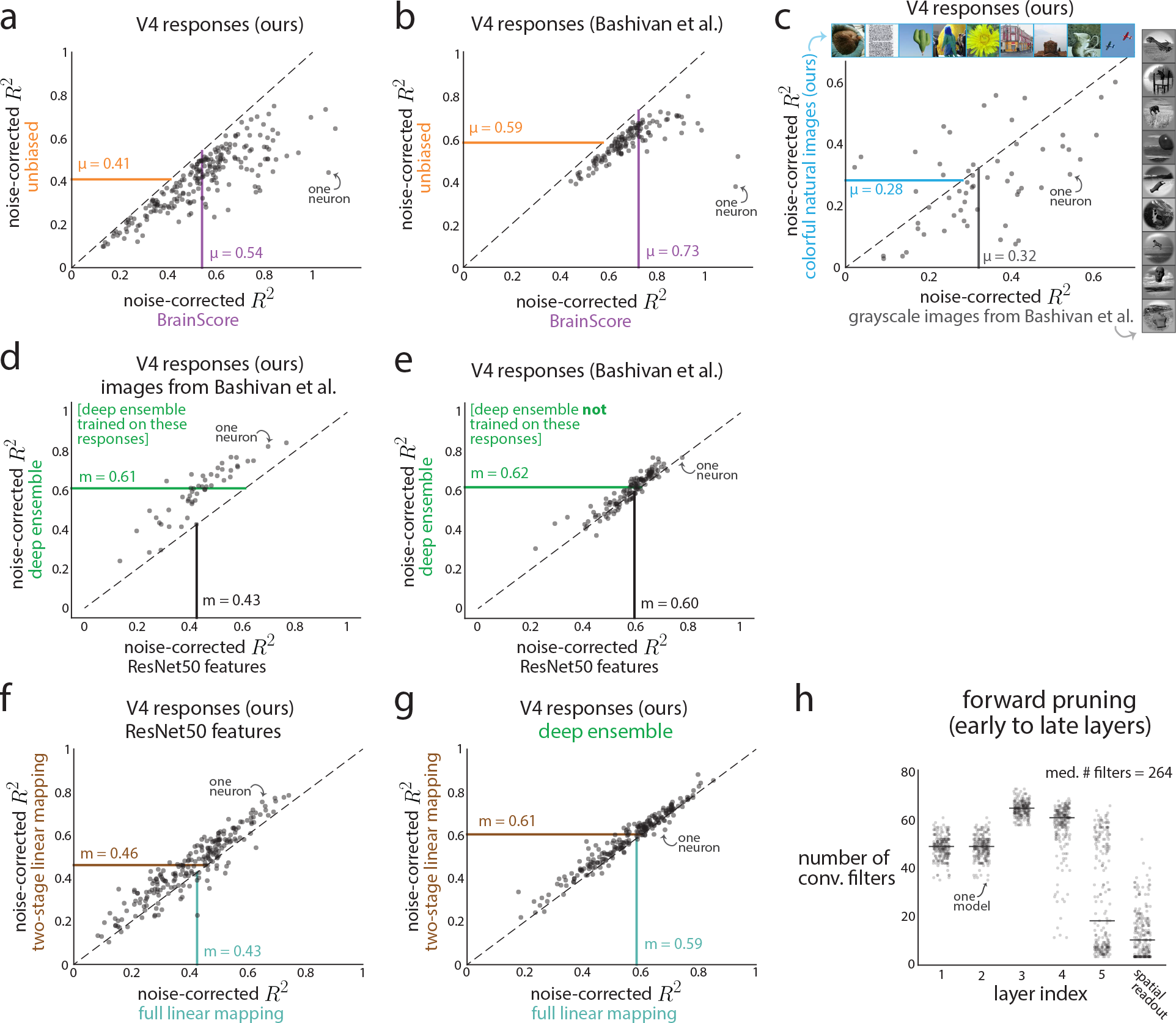
Comparing our noise-corrected *R*^2^ values versus those reported in a previous study. Comparing noise-corrected *R*^2^s across studies can be difficult, as they often differ in spike sorting procedures, the types and numbers of images presented, how the noise-corrected *R*^2^ metric is computed, and how DNN features are mapped to V4 responses. Indeed, for linearly mapping task-driven DNN features to V4 responses, we find a difference between our reported noise-corrected *R*^2^ = 0.45 (Fig. 1**b**) with another reported noise-corrected *R*^2^ = 0.89 from a previous study [40]. Here, we determine that this difference is primarily due to different noise-corrected *R*^2^ metrics and spike sorting criteria. **a**. A key difference between our study and [40] is our noise-corrected *R*^2^ metric. [40] use the BrainScore *R*^2^, which computes an *R*^2^ ceiling by splitting repeats into two halves, corrected with the Spearman-Brown procedure [30]. The BrainScore *R*^2^ is computed with the following pseudocode: 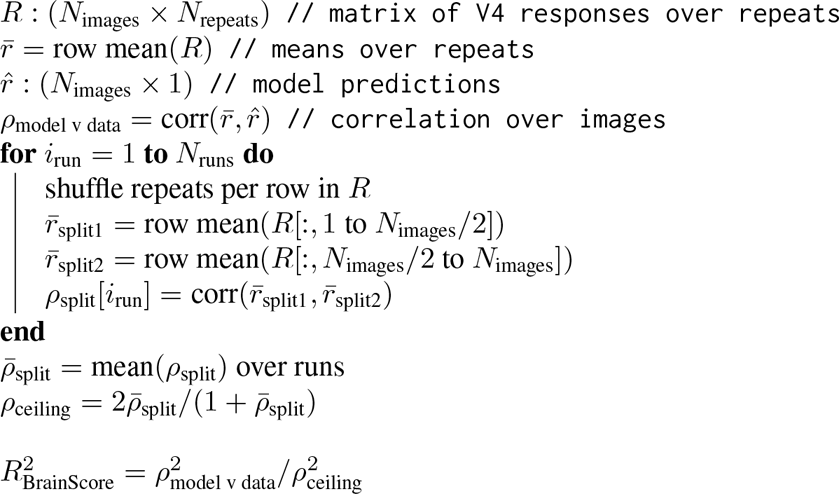 We use a recently-proposed unbiased noise-corrected *R*^2^ metric (mathematically defined in Methods), which was shown to be more consistent at estimating the true *R*^2^ versus other proposed *R*^2^ metrics [106]. For each of our 219 recorded neurons on 4 held out sessions, we computed the unbiased *R*^2^ versus the BrainScore *R*^2^ using a linear mapping (ridge regression) between ResNet50 features and V4 responses (cross-validated). We noticed a sizable increase of the BrainScore *R*^2^ (dots to the right of dashed line, ∆*R*^2^ *≈* 0.15), including some neurons with BrainScore *R*^2^ *>* 1 (dots to the right of 1), likely caused by too few repeats to properly estimate *R*^2^. This overestimation in *R*^2^ was expected and motivation for an unbiased *R*^2^ [31]. This increase appeared to be a shift for every neuron, as the unbiased *R*^2^ was correlated with the BrainScore *R*^2^ across neurons (*ρ* = 0.84). Each dot denotes a V4 neuron; *µ* denotes the mean *R*^2^ across neurons. **b**. Because [40] uses the BrainScore metric, we wondered if this entirely explained the difference between the reported *R*^2^s. We re-computed the BrainScore and unbiased *R*^2^ metrics for the publicly-available V4 responses from [40] with ResNet50 features and found a similar difference in mean noise-corrected *R*^2^ with a larger Brain-Score *R*^2^ (dots to the right of dashed line, ∆*R*^2^ *≈* 0.15). Again, this increase appeared to be a shift, as the *R*^2^s were correlated across neurons (*ρ* = 0.86). We note that our reported BrainScore *R*^2^ = 0.73 was lower than that reported by [40] (*R*^2^ = 0.89) for the same data. To account for this difference, we trained a factorized linear mapping [110] (same as used in [40]) and achieved a BrainScore *R*^2^ = 0.83; we suspect using their retinal transformation would then achieve the same BrainScore *R*^2^. We conclude that the BrainScore *R*^2^ metric adds ∼ 0.15 to an unbiased *R*^2^ metric; however, for the same unbiased *R*^2^ metric, we see a difference of ∆*R*^2^ *≈* 0.2 (compare orange *µ* between **a** and **b**; similar for BrainScore *R*^2^). Thus, a difference in metric contributes to differences in reported *R*^2^s but does not provide a full explanation. **c**. We checked another possibility—in our study, we presented colorful natural images while [40] presented grayscale images with foreground objects placed on unrelated natural backgrounds (see inset images). To see if this difference in image type could lead to differences in *R*^2^, we used one recording session to present 600 colorful natural images and 600 images from [40]. We found that when predicting these V4 responses with ResNet50 features, there was only a modest increase in *R*^2^ for the grayscale images (mean *R*^2^ = 0.32 for grayscale images versus mean *R*^2^ = 0.28 for colorful images). We also observed that these *R*^2^s were correlated across neurons (*ρ* = 0.39). Thus, the differences in image statistics likely was not a contributing factor in the differences in unbiased *R*^2^ between our responses and responses in [40]. We conclude that this difference in *R*^2^ arises because of differences in spike sorting and electrode unit criteria. [40] have stricter criteria for retaining recorded units (neurons must be present across multiple sessions, single-unit isolation, etc.), whereas our criteria are less strict (neurons need only be present for one session, multi-unit activity is possible). This highlights the need to test one’s model on multiple data sets from different laboratories [30]. **d**-**e**. Compared to a task-driven DNN model that appears to generalize well for any V4 neuron, we tailored our data-driven deep ensemble model to predict only our recorded V4 responses. It was unclear if our deep ensemble model could generalize to V4 neurons on which it was not trained (i.e., generalize to “out-of-distribution” V4 neurons). To test this, we sought to use the deep ensemble model to predict the V4 responses from [40]. To make a fair comparison, we recorded V4 responses to images from [40]; we confirmed that our deep ensemble model predicted responses to these images (**d**, median unbiased noise-corrected *R*^2^ m = 0.61) to the same extent as those to natural images (Fig.1**b**, median *R*^2^ m = 0.61). Similarly, using ResNet50 features had nearly identical prediction performances (**d**, median unbiased *R*^2^ m = 0.43; Fig. 1**b**, median *R*^2^ m = 0.43). This suggests that our deep ensemble model could predict our V4 responses to images from [40] to the same extent as those to colorful natural images, consistent with ResNet50’s predictions (**c**). Next, we performed the same analysis but for V4 responses from [40]. Using ResNet50 features increased prediction performance (**d**, m = 0.43 to **e**, m = 0.60), consistent with the increased prediction performance between our V4 responses to colorful natural images (**a**, orange, *µ* = 0.41) and V4 responses from [40] (**b**, orange, *µ* = 0.59). We expected prediction performance for the deep ensemble to worsen between our V4 responses and responses from [40], as our deep ensemble model was optimized only for our recorded V4 responses. However, this was not the case: Prediction performance remained at the same level between the two (**d**, m = 0.61 versus **e**, m = 0.62). This suggests that our deep ensemble model can generalize to new V4 neurons, although we lose the performance gains from training on these V4 neurons. We expect that as our deep ensemble model is trained on responses from more V4 neurons, its generalization ability will increase, surpassing that of ResNet50. **f**-**g**. For completeness, we also consider a newly proposed mapping between DNN features and V4 responses that factorizes a linear mapping into a ‘spatial’ stage and a channel ‘mixing’ stage [110] (see Methods). We found a modest increase in prediction performance for this factorized linear mapping versus a linear mapping identified with ridge regression, both when using ResNet50 features (**f**, median unbiased *R*^2^ m = 0.46 versus m = 0.43) and the output embeddings of the deep ensemble model (**g**, median unbiased *R*^2^ m = 0.61 versus m = 0.59) for V4 responses to natural images in our 4 held out recording sessions. We suspect that this increase, smaller than those observed previously [40, 110], arises because we have roughly double the number of images shown per session on which to train and test the mappings. **h**. We designed a procedure to prune filters that, if ablated, led to little change in the model’s output. Our pruning procedure starts with filters in the deepest layers and proceeds backwards to filters in the earliest layers. After pruning, we found the earliest layers had larger numbers of filters ( ∼ 50 filters each, Fig. 3**a**) than the deepest layers (5-10 filters each, Fig. 3**a**). However, we wondered if this trend arose because we pruned filters in the deepest layer first. To test for this, we reversed the order of pruning to begin with the earliest layer (i.e., layer 1) and continue to the deeper layers. This reversal led to a larger number of filters overall (median number of filters across neurons was 264 filters versus the 164 filters in the deep-to-early-layer pruning), yet we still found the same trend of a larger number of filters in early versus deeper layers (compare **h** with Fig. 3**a**). We conclude that the consolidation step identified in our compact models was not due to the way in which the model was pruned.

**Extended Data Figure 3:**
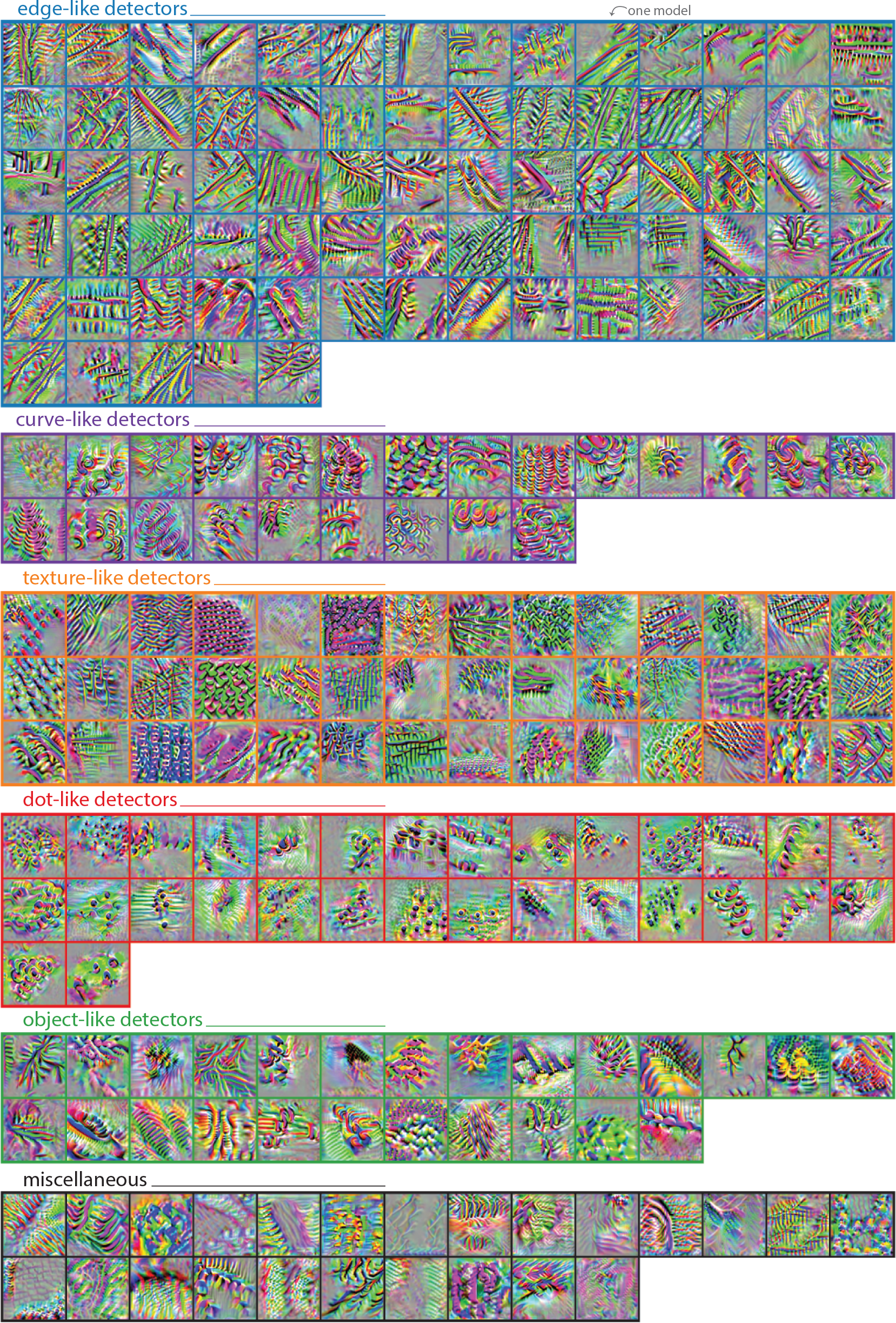
Maximizing synthesized images for all compact models. For each compact model, we computed the maximizing synthesized image via gradent ascent techniques (Fig. 2**a**). Here, we show these images for all compact models (one image per compact model). For visual clarity, we loosely grouped images into categories by eye. We make no claims about grouping in V4 neurons; in fact, the mean signal correlation squared across all pairs of compact models was low (*ρ*^2^ = 0.11 between model responses to 10,000 normal images). This remained true when controlling for spatial receptive field location (Fig. 3**c**, layer 5). Thus, the stimulus preferences across V4 neurons appear to be largely heterogeneous, allowing for a highly-expressive set of features for downstream processing (e.g., IT neurons).

**Extended Data Figure 4:**
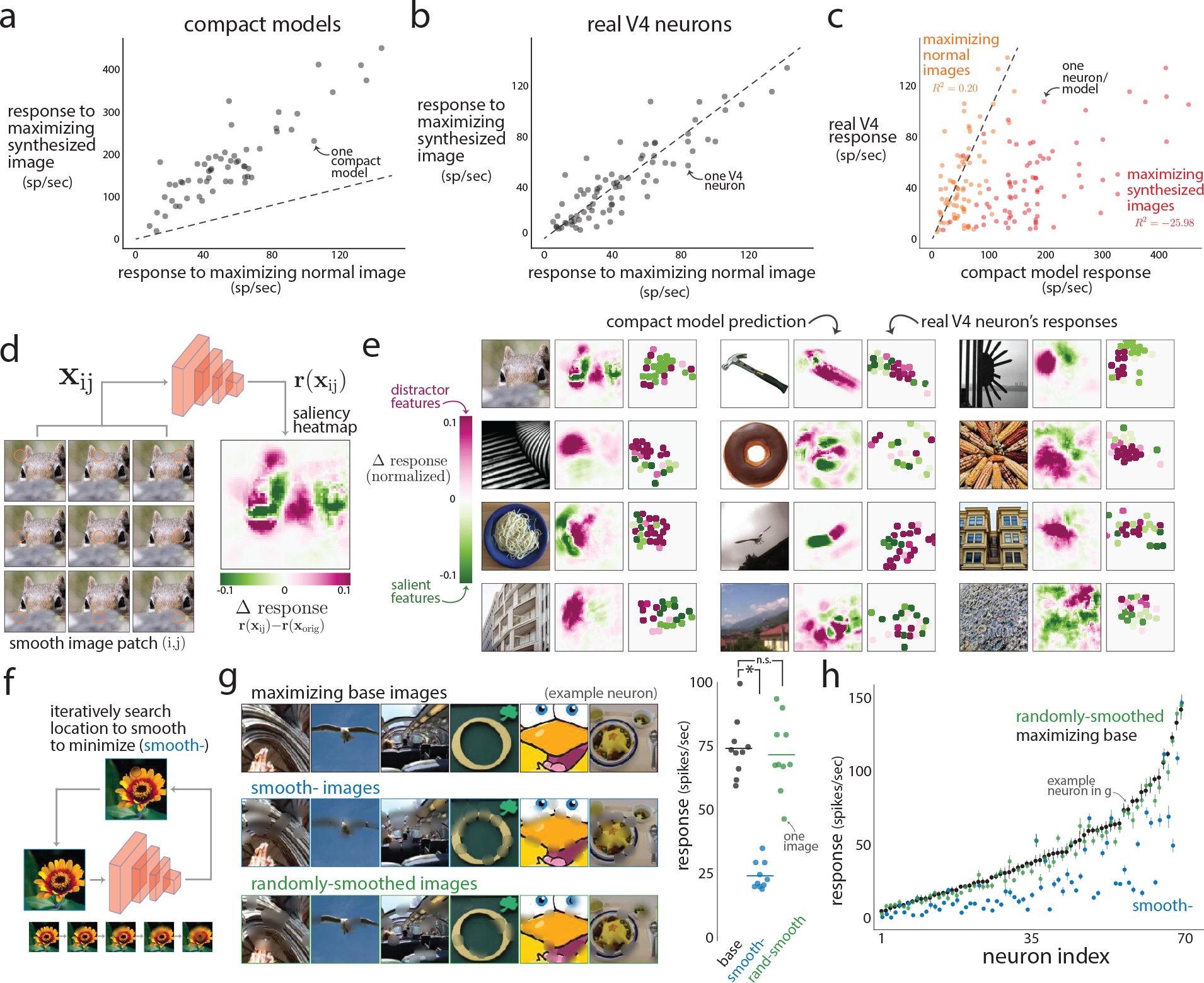
Further causal tests of the compact models. Our compact models’ predictions held up to causal testing (Fig. 2), including identifying maximizing normal and synthesized images. Here, we compare which type of these images better drives V4 responses. In addition, we present results of further causal tests, including identifying saliency maps and “adversarial smooth” images. **a**-**c**. We found that the compact models predicted much stronger V4 responses to maximizing synthesized images versus maximizing (normal) images (**a**, dots above dashed line; difference of means between maximizing synthesized and maximizing normal: 111.7 spikes/sec, *p <* 0.002, permutation test). However, this was not the case for the real responses—both types of images evoked similarly large responses (**b**, dots hug dashed line, difference of means not significant, *p* = 0.836, permutation test). This difference largely stems from the compact models’ inability to predict V4 responses to maximizing synthesized images (**c**, red dots, coefficient of determination *R*^2^ = *−* 26.0, not noise-corrected) whereas their prediction for maximizing normal images remains relatively intact (**c**, orange dots, coefficient of determination *R*^2^ = 0.2, not noise-corrected). Each dot denotes a neuron’s response, averaged over repeats and maximizing images (if multiple maximizing images were shown for a V4 neuron, see Fig. 2**b** and **c**). This inability to predict maximizing synthesized images was not unexpected—the optimizing procedure had full access to every weight and every pixel to optimize an image customized for that compact model. Moreover, the resulting synthesized images were well outside the distribution of training images, and we would expect poor prediction for these outlying regions of image space. This was one motivation for training our deep ensemble model with closed-loop active learning (Fig. 1**f**) in which we trained the model on out-of-distribution images. We were also suprised that the maximizing normal images evoked V4 responses as large as those to maximizing synthesized images (**b**). This suggests that one cannot rule out choosing from a large pool of candidate images (in this case, 500,000 candidate images) to maximally drive V4 responses. **d**. A commonly-used approach to explain a DNN’s output is to identify which parts of an input image are the most relevant or “salient” for the DNN’s prediction; this approach is called saliency analysis [123]. One implementation is to smooth a small patch of the image (small orange circles in example images denote smoothed patches; these circles were not present in actual stimuli) and see if the resulting response is larger or smaller than the response to the original image. An increase in response (pink) indicates that the visual feature within the smoothed patch is distracting, as removing this feature leads to a *larger* response. Likewise, a decrease in response (green) indicates a salient or excitatory visual feature. We passed as input a set of images where the (*i, j*)th image had a smoothed image patch centered at (*i, j*). We then formed a heatmap of the resulting responses **r** based on the (*i, j*) of the image patch (‘saliency heatmap’). For this example image of a squirrel and the chosen compact model, the most salient features are the eyes (green regions), while the most distracting features are the fur texture and the edges around the left eye (pink regions). **e**. In our causal experiments, we probed the trained compact models to identify maximizing normal images (Fig. 2**b**). For each maximizing normal image, we computed the saliency heatmap of the compact model (‘compact model prediction’) following the procedure in **d**. We then used these predictions to smooth 25 non-overlapping image patches that led to the largest changes in responses (where the number 25 was chosen as a compromise between covering as much as the image as possible within recording time constraints). On a following session, we showed each ‘base’ image as well as the 25 images, each with one smoothed image patch. For each example image shown here, we matched a V4 neuron with the image’s corresponding compact model (in the same way as in Fig. 2) and computed the resulting saliency heatmap for V4 neurons (rightmost panels). Responses were z-scored using the mean and standard deviation estimated with the V4 neuron’s responses to all normal images shown in the session. We found that V4 neurons did vary their responses to local smoothing and that these changes in responses largely matched those predicted by the compact models. Thus, for a given image, a V4 neuron’s response can be suppressed and excited by different local visual features; our compact models can be used to predict which features at which locations. **f**. Inspired by the saliency approach in **d** and **e**, we causally tested our compact models by having them predict which visual features of an image to smooth in order to minimize a V4 neuron’s response. We first began with the compact model’s maximizing normal image as a base image. Then, in a greedy manner, we iteratively chose an image patch to smooth that led to the smallest response as predicted by the compact model (‘smooth-’, see orange circle). Successive iterations added image patches that did not overlap with previously-chosen image patches. This led to an image with specific visual features smoothed away. Bottom inset: a sequence of images for which a base image is cumulatively smoothed at different patches determined by the compact model’s predictions; the final smoothed image is the rightmost. **g**. Example base images (left, top row, ‘maximizing base images’), smoothed versions to minimize the model output response as predicted by a compact model (left, middle row, ‘smooth-images’), and images for which randomly-chosen patches were smoothed as a control (left, bottom row, ‘randomly-smoothed images’). The randomly-smoothed and smooth-images had the same number of pixels smoothed. These example maximizing base images and randomly-smoothed images elicited similarly large responses from a V4 neuron (right, black versus green dots, *p* = 0.70, permutation test) whereas the smooth-image led to a substantially smaller response (black versus blue dots, *p <* 0.002, permutation test, asterisk). Each dot denotes the repeat-averaged response to one image. **h**. Responses for all V4 neurons from two recording sessions (each session from a different animal). V4 neurons were matched to compact models via held out images (same procedure as used in Fig. 2**b**-**f**). For one session, only one base image was shown per compact model; for these images, dots denote repeat-averaged responses with no error bars. For the other session, we presented 10 base images (and their smooth- and randomly-smoothed counterparts) per neuron. For this session, dots denote the average response over the 10 images (and their repeats), and error bars denote s.e.m. over the 10 images. Neurons were sorted based on mean response to the base images. We found that responses to smooth-images were roughly half as small as responses to the base images across V4 neurons (blue versus black dots, normalized percent change computed as 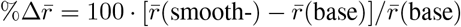 : mean 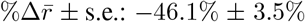). Little to no decrease between maximizing base and randomly-smoothed images (green versus black dots, mean 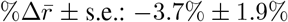). Thus, in a causal test, the compact models accurately predicted which visual features (and their spatial information) were most salient to the V4 neurons. This provides further evidence that the compact models accurately capture the stimulus preferences of V4 neurons— and what visual features in those preferred stimuli are most important to the V4 neuron.

**Extended Data Figure 5:**
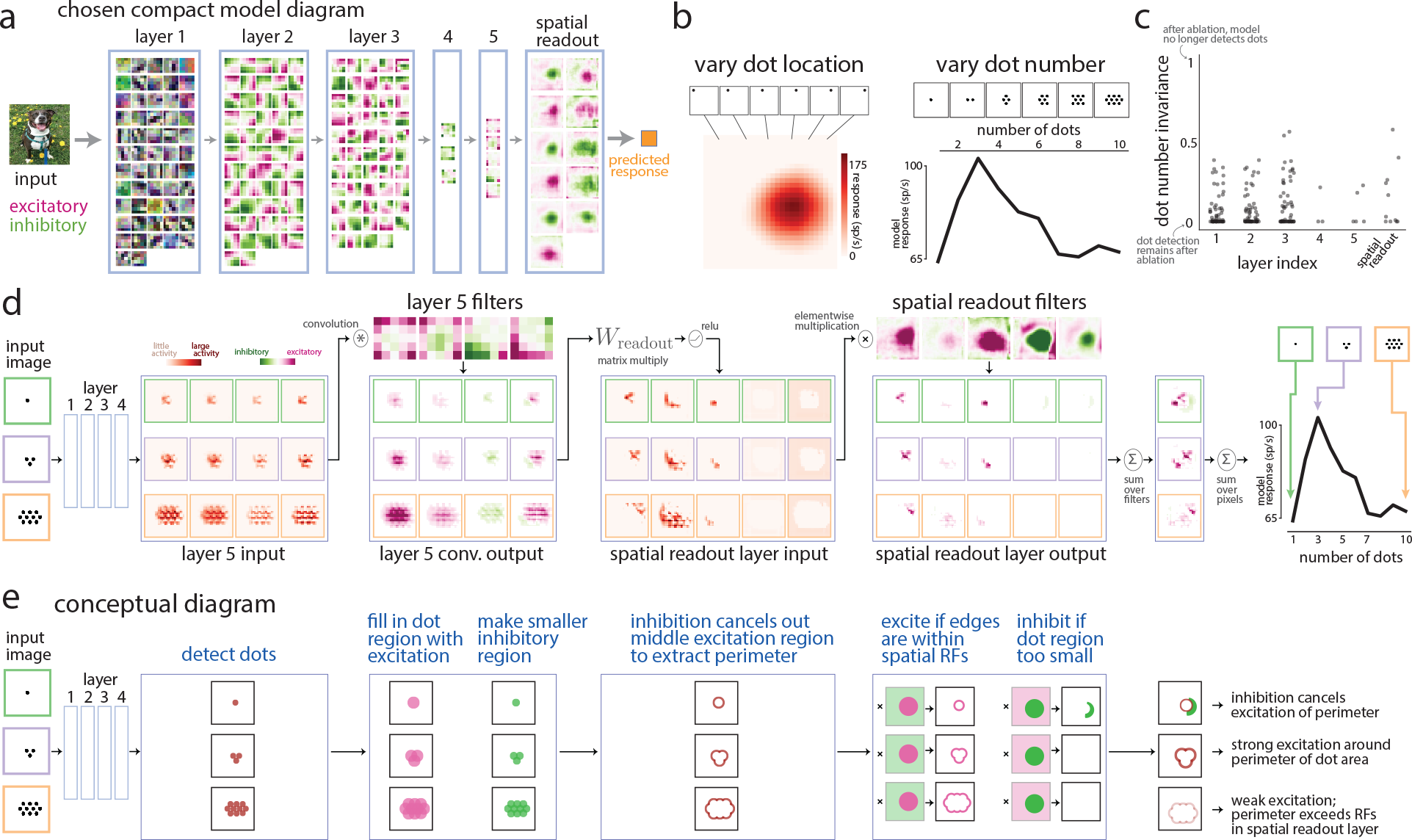
Explaining a dot-detecting compact model’s selectivity for multiple dots. To understand the computations within a compact model, we chose to investigate a particular compact model that resembled a dot detector (Fig. 4). We focused on the model’s selectivity to dot size (Fig. 4**b**) and uncovered a simple computation by isolating the filters that contributed to dot size selectivity (Fig. 4**c**-**h**). However, we suspected this compact model was also selective to the number of dots, as its preferred stimuli typically had two to five dots in the image (Fig. 4**a**). Here, we investigate this model’s dot number selectivity. **a**. Diagram of the compact model that resembles a dot detector. We expected to see dot-like filters (i.e., middle excitation surrounded by inhibition or vice versa) in layers 1-3 but found none. This led us to identify corner and large edge detecting filters that ultimately contributed to dot size selectivity (Fig. 4**c**-**h**). **b**. Besides varying dot size, we also varied the location of a small dot with a radius of 5 pixels (left, ‘vary dot location’) and dot number (right, ‘vary dot number’). The compact model (same as in **a** and Fig. 4) preferred three dots to the center right (with dot radii of 5 pixels, from Fig. 4**b**). Given this compact model’s preferred stimuli (Fig. 4**a**) and selectivity to dot location (left), dot number (right), and dot size (Fig. 4**b**), we conclude that this compact model is a dot detector. **c**. In the same manner as we identified filters contributing to dot size selectivity (Fig. 4**c** and **d**), we ablated each filter and measured the model’s dot number invariance. A dot number invariance of 1 indicates that after ablating a filter, the model is no longer able to detect different numbers of dots; a dot number invariance of 0 indicates no change in the model’s output (i.e., dot number selectivity remains intact). We found that none of the filters contributed strongly to dot number selectivity (i.e., no filter, once ablated, led to a large increase in dot number invariance); this differs from the highly contributing filters for dot size selectivity (compare **c** with Fig. 4**d**, filters with DSI > 0.75). This indicates that dot number selectivity emerges from the last spatial readout layer. **d**. To understand the specific computations of dot number selectivity, we passed in three input images with different numbers of dots (‘input image’). We then observed the resulting activity and filter weights for layer 5 and the spatial readout layer. We noticed that the input to layer 5 (‘layer 5 input’) appeared to detect the presence of a dot at any given location (matching our intuition for identifying a single dot in Fig. 4**f**-**h**). The layer 5 filters (chosen as those with the largest dot number invariance in **c**) appeared to extract specific patterns of the detected dots: After convolution, some filters formed large excitatory regions around the dots whereas others formed smaller regions of inhibition (‘layer 5 conv. output’). After passing through a linear combination of its filters (by multiplying with readout matrix *W*_readout_) and the ReLU activation function, we found that most of the dot regions were extinguished with some filters activated by the edges or shells of the dot region (‘spatial readout layer input’). This activity was then excited or suppressed by spatial readout filters that act as spatial receptive fields. After taking a linear combination over filters (‘sum over filters’), we observed that a single small dot had both excitation and inhibition (top activity map), while three dots led to large activity around the region of the dots (middle activity map). Many dots led to little to no activity (bottom activity map). Summing over spatial information (i.e., pixels) recreated the selectivity to dot number (rightmost plot). We illustrate the circuit mechanisms in **d** with a conceptual diagram. The key concept is that the model identifies a region of dots, extracts the shell around this region (by creating a larger excitatory region and a smaller inhibitory region), and queries the size of this shell. Too small a region (i.e., a single dot) leads to weak activity and inhibition (rightmost, top activity map). Too large a region leads to weak excitation, as the shell of the region is outside the spatial receptive field. Only an appropriately sized region (i.e., a certain number of dots) will fit within the spatial receptive field, leading to strong excitation. This results suggest that a key computation in dot number selectivity is to extract “regions” of interest and then to identify the edges of these regions. If the number of dots is too large, the region’s edge will be outside the spatial receptive field, yielding a small response. This may be a reoccurring theme in the visual cortex for other visual features.

**Extended Data Figure 6:**
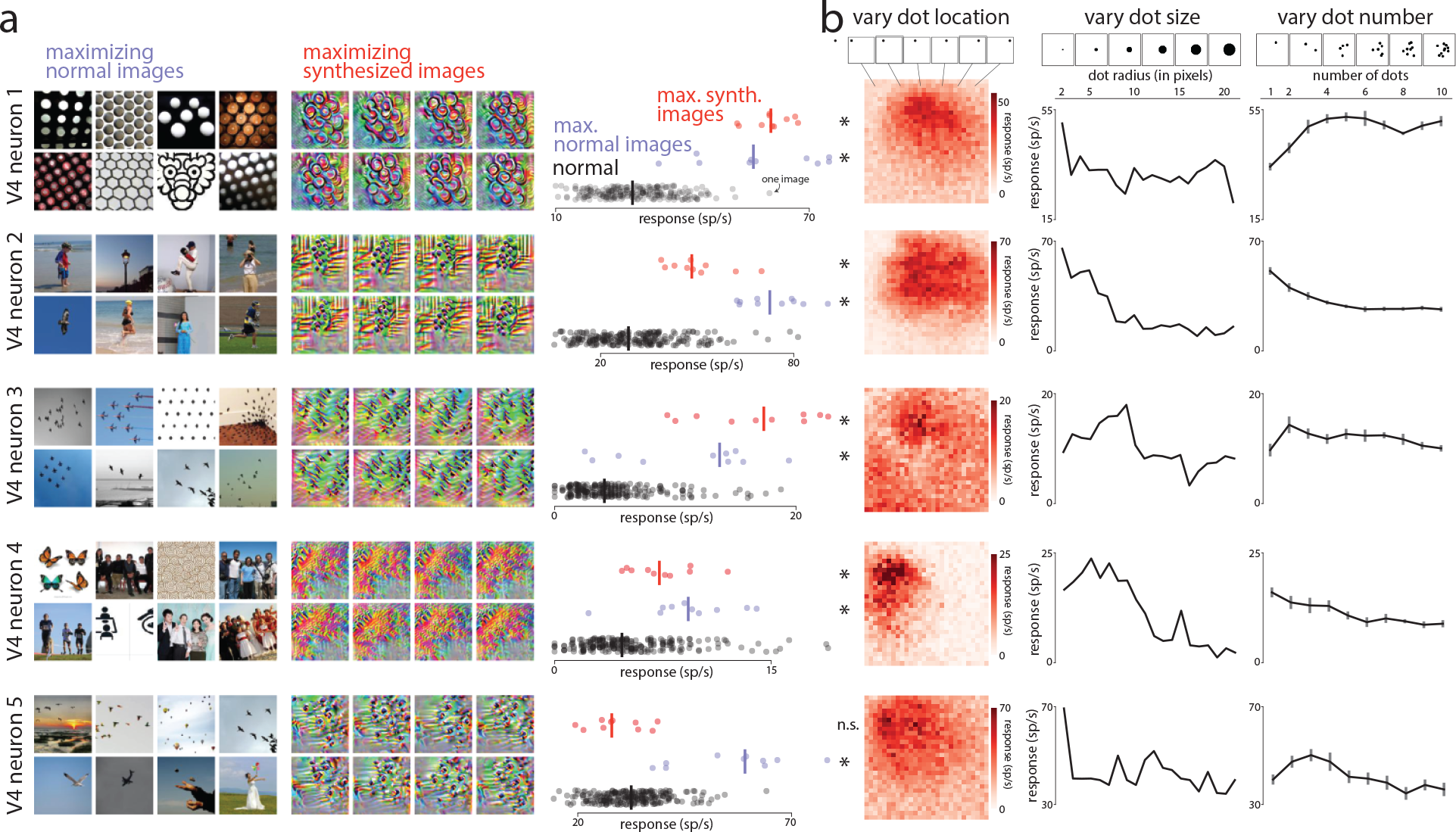
Identifying real V4 dot detectors with causal tests. To confirm the presence of dot-detecting V4 neurons, we ran causal tests specifically tailored for compact models that resemble dot detectors. We first identified compact models by training on previous recording sessions. From the identified compact models, we chose 5 compact models that most resembled dot detectors based on their stimulus preferences (i.e., maximizing normal and synthesized images) and their responses to artificial dot stimuli (see Fig. 4**a** and **b** and Ext. Data Fig. 5). We note that the chosen dot detecting compact model in Figure 4 was not one of the chosen, as this compact model matched to a V4 neuron from another animal. For a future recording session, we presented the maximizing normal and synthesized images of the 5 chosen models as well as artificial dot stimuli (same dot stimuli as in Fig. 4**b** and Ext. Data Fig. 5**b**). We identified the 5 recorded V4 neurons that best matched the predictions of the 5 compact models (by computing the noise-corrected *R*^2^ from all other images shown in the session, same procedure as in Fig. 2**b**-**f**) and show their responses here. **a**. The maximizing normal images (left, examples) and maximizing synthesized images (middle, examples) chosen from the 5 compact models tended to more strongly drive V4 responses than responses to normal images (right, ‘max. synth. images’ and ‘max. images’ dots more to the right than ‘normal’ black dots). Each dot is the repeat-averaged response to one image; lines denote medians. All maximizing stimuli yielded median responses significantly greater than the median response to normal images (*p <* 0.02, one-sided permutation test, asterisks) except one set of maximizing synthesized stimuli (bottom row, V4 neuron 5, red dots, *p* = 0.922, one-sided permutation test). This failure could be from instability in the electrode array (i.e., the V4 neuron on which the compact model was trained no longer was accessible by the electrode array) or due to model mismatch. **b**. V4 responses to the artificial dot stimuli that varied in dot location (left), dot size (middle) and number of dots (right). Same format as in Figure 4**b** and Extended Data Figure 5. Dot locations were subsampled to 28 *×* 28 locations to limit the number of images. Error bars in ‘vary dot number’ denote 1 s.e.m. across 10 different images, where each image had the same number of dots but in different locations randomly chosen to be nearby the preferred location (see Methods). We found that these V4 neurons had preferred dot locations (**b**, left), preferred dot sizes (**b**, middle), and preferred numbers of dots (**b**, right), consistent with these V4 neurons being dot detectors. Thus, these results confirm the presence of dot detectors in V4. We observed diverse selectivity to dot size, including V4 neurons selective to the tiniest dots (neurons 1, 2, and 5) and small dots (neurons 3 and 4). Similarly, we observed selectivity to one dot (neurons 2 and 4), 2 dots (neuron 3), and 3 or more dots (neurons 1 and 5). Thus, even within the class of dot detectors, there appears to be large diversity in stimulus preferences.

**Extended Data Figure 7:**
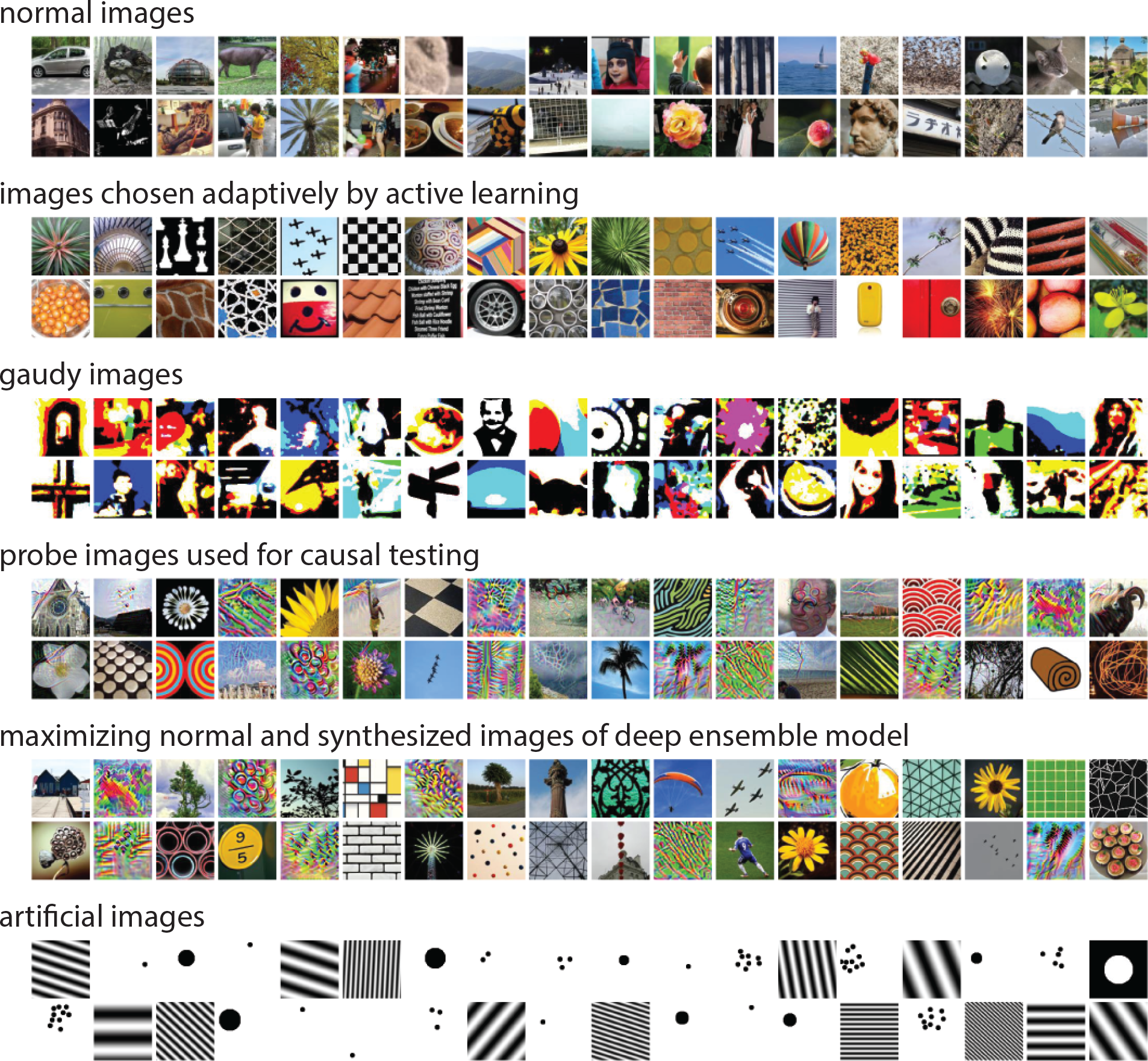
Example visual stimuli shown in experiments. Example images for each type of visual stimuli shown in our experiments. Example images were randomly selected for each type. See Methods for details about how images were chosen or generated.

## Extended Data Tables

**Extended Data Table 1:**
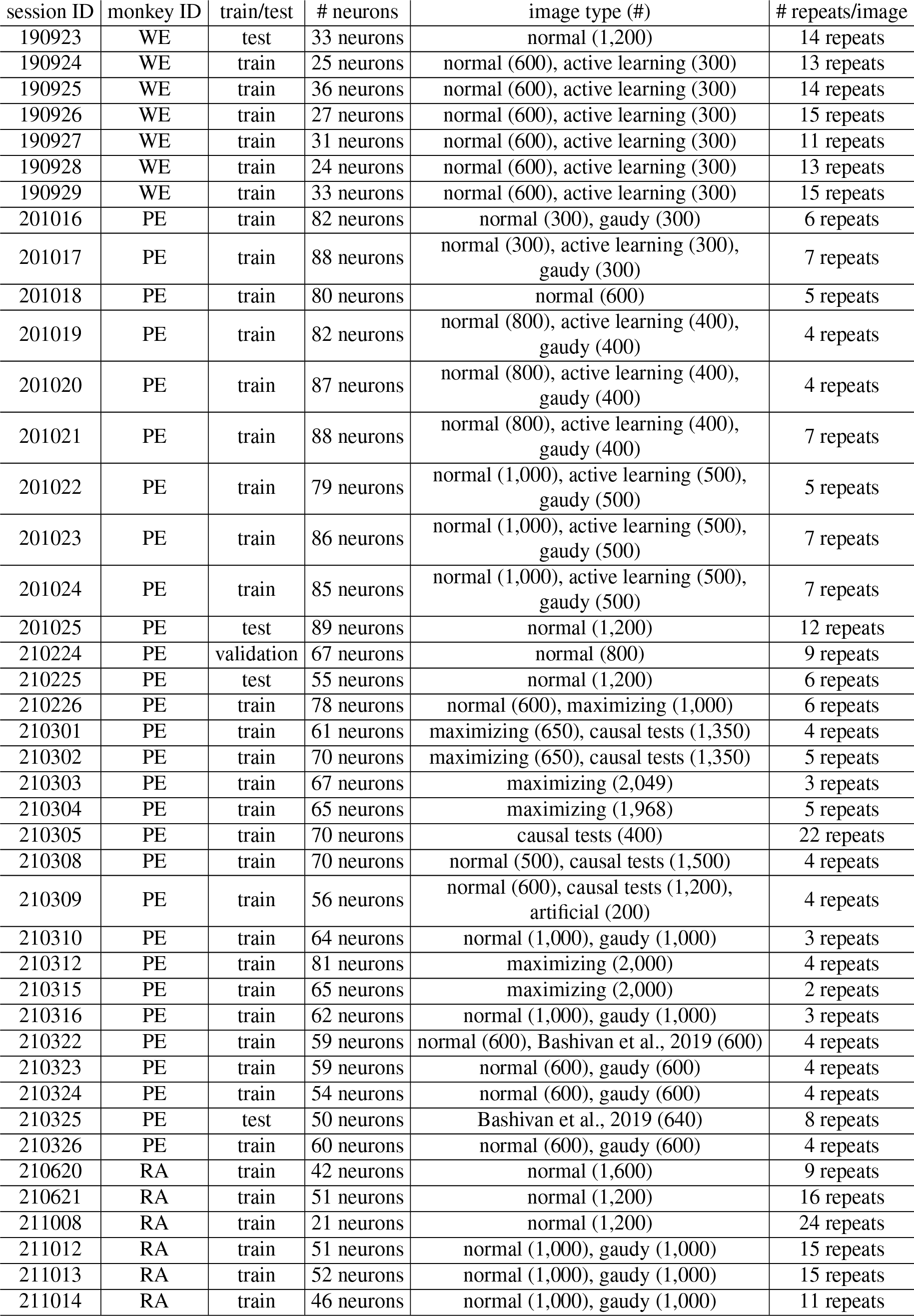
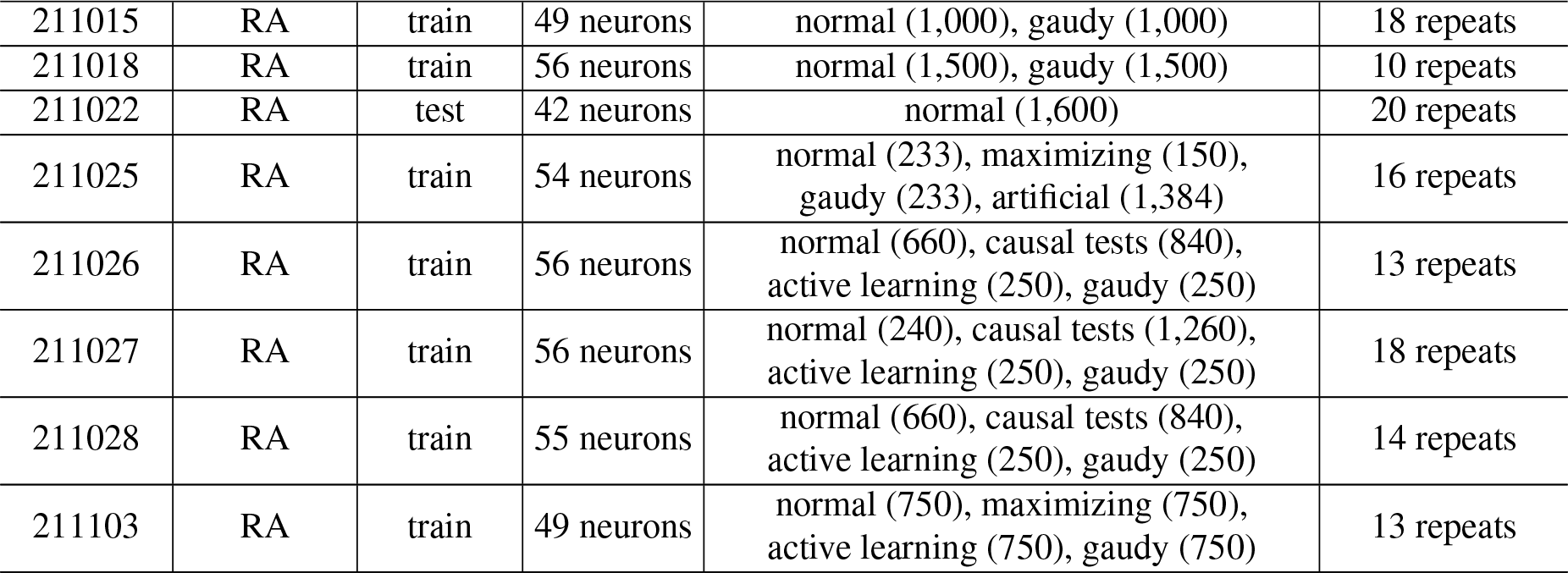
Details of recording sessions and visual stimuli. Details for all 50 recording sessions from 3 monkeys (WE, PE, RA). Session ID is the date collected (YY/MM/DD). Sessions were either used for training (‘train’, 45 sessions), testing (‘test’, 4 sessions), or validation (‘validation’, 1 session). We re-used some of causal experiment sessions (e.g., session 211026) as training sessions for the deep ensemble model; the probe images for these sessions were generated from compact models trained only on preceding sessions (e.g., sessions 190923-211025). The reported number of neurons is after removing units with low SNR, low firing rates, etc. (see Methods). We also report the visual stimuli types and numbers. For sessions 210322 and 210325, we presented images from a dataset released by [40] for comparison (Ext. Data Fig. 2). For a few sessions, we presented maximizing synthesized images not only for the output of the compact models but also for internal filters of the compact models; these images were used for exploratory analyses. We also report the median number of repeats across images; the number of repeats may vary across images due to recording session length and the number of correct/incorrect trials performed by the animal.

## Notes

### Competing Interest Statement

The authors have declared no competing interest.

